# Deciphering the ferroptosis pathways in dorsal root ganglia of Friedreich ataxia models. The role of LKB1/AMPK, KEAP1, and GSK3β in the impairment of the NRF2 response

**DOI:** 10.1101/2024.05.10.593481

**Authors:** Arabela Sanz-Alcázar, Marta Portillo-Carrasquer, Fabien Delaspre, Maria Pazos-Gil, Jordi Tamarit, Joaquim Ros, Elisa Cabiscol

**Affiliations:** Departament de Ciències Mèdiques Bàsiques, IRBLleida, Universitat de Lleida, Catalonia, Spain

**Keywords:** Friedreich ataxia, frataxin, dorsal root ganglia, ferroptosis, NRF2, LKB1/AMPK pathway

## Abstract

Friedreich ataxia (FA) is a rare neurodegenerative disease caused by decreased levels of the mitochondrial protein frataxin. Frataxin has been related in iron homeostasis, energy metabolism, and oxidative stress. Ferroptosis has recently been shown to be involved in FA cellular degeneration; however, its role in dorsal root ganglion (DRG) sensory neurons, the cells that are affected the most and the earliest, is mostly unknown. In this study, we used primary cultures of frataxin-deficient DRG neurons as well as DRG from the FXN^I151F^ mouse model to study ferroptosis and its regulatory pathways. A lack of frataxin induced upregulation of transferrin receptor 1 and decreased ferritin and mitochondrial iron accumulation, a source of oxidative stress. However, there was impaired activation of NRF2, a key transcription factor involved in the antioxidant response pathway. Decreased total and nuclear NRF2 explains the downregulation of both SLC7A11 (a member of the system Xc, which transports cystine required for glutathione synthesis) and glutathione peroxidase 4, responsible for increased lipid peroxidation, the main markers of ferroptosis. Such dysregulation could be due to the increase in KEAP1 and pGSK3β (Tyr216), which promote cytosolic localization and degradation of NRF2. Moreover, there was a deficiency in the LKB1/AMPK pathway, which would also impair NRF2 activity. AMPK acts as a positive regulator of NRF2 and it is activated by the upstream kinase LKB1. The levels of LKB1 were reduced when frataxin decreased, in agreement with reduced pAMPK (Thr172), the active form of AMPK. SIRT1, a known activator of LKB1, was also reduced when frataxin decreased. In conclusion, this study demonstrated that frataxin deficiency in DRG neurons disrupts iron homeostasis and the intricate regulation of molecular pathways affecting NRF2 activation and the cellular response to oxidative stress, leading to ferroptosis.

## 1. Introduction

Friedreich ataxia (FA) is a rare hereditary neurodegenerative disease caused by mutations in the frataxin gene (*FXN*), resulting in a reduction in the frataxin content. In most patients, frataxin deficiency is caused by the presence of GAA-triplet expansion in homozygosity in the first intron of the *FXN* gene [1]. Around 4% of patients are compound heterozygotes: They present GAA repeats in one allele and point mutations or deletions in the other [2]. The disease is characterized by progressive limb and gait ataxia, limb muscle weakness, scoliosis, diabetes, and hypertrophic cardiomyopathy [3], with the latter being the main cause of death in patients with FA [4] The symptoms usually start in childhood or adolescence, but they can appear later [5].

Frataxin is a mitochondrial protein, and its deficiency leads to mitochondrial dysfunction, with iron accumulation and oxidative stress, denoted by uncontrolled production of reactive oxygen species (ROS). Although it is generally accepted that frataxin activates iron–sulfur biogenesis, iron–sulfur deficiency is not a universal consequence of frataxin deficiency, suggesting that the role of frataxin in this process is not essential [6]. Iron deposits or accumulation have been observed in frataxin-deficient models, but their role in disease progression remains unclear [7] [8] It has also been suggested that iron accumulation may be due to constitutive activation of the iron deficiency response pathway, leading to decreased expression of key metabolic enzymes and mitochondrial complex components [9].

Although much research has been conducted, there is no effective cure for FA. The first pathological changes occur in the dorsal root ganglia (DRG) with the loss of large sensory neurons, followed by degeneration of the spinocerebellar and corticospinal tracts, and atrophy of the large sensory fibers in peripheral nerves [10]. DRG neurons express the highest levels of frataxin and display high vulnerability to frataxin downregulation. For this reason, the consequences of frataxin depletion have been studied in DRG, at the histological level, using conditional knockout mice [11] [12] and samples from patients with FA [13] [14]. By using primary cultures of frataxin-deficient DRG neurons, our group has shown alterations in several parameters, including decreased mitochondrial membrane potential, altered calcium homeostasis, and mitochondrial pore opening [15][16] [17] In a recent report, we described mitochondrial iron accumulation and deficient mitochondrial respiration, which resulted in a decreased NAD^+^/NADH ratio and, consequently, a decline in sirtuin activity. There were elevated levels of the acetylated (inactive) form of mitochondrial superoxide dismutase 2 (SOD2), a SIRT3 substrate, contributing to oxidative stress [18].

The central role of iron dysfunction and oxidative stress in FA has been highlighted by several reports indicating that frataxin-deficient cells are hypersensitive to erastin, a drug that causes glutathione depletion and is a well-known inducer of ferroptosis [19] [20]. Ferroptosis has been defined as a regulated form of cell death triggered by the accumulation of lipid peroxidation products in membranes, and determined by an iron-dependent ROS overload [21]. At the morphological level, cells display a reduction in mitochondrial cristae, but the cell membrane does not break, the nucleus remains at its normal size, and there is no chromatin condensation [22]. Ferroptosis requires the presence of redox-active iron and the reduction of cellular antioxidant defenses via nuclear factor erythroid 2–related factor 2 (NRF2), a transcription factor involved in adapting to oxidative stress conditions [23]

While mitochondrial iron accumulation, increased oxidative stress, lipid peroxidation, and NRF2 impairment are all found in various frataxin-deficient models, are hallmarks of ferroptosis, this process has only recently been implicated in the disease [19] [24]. Interestingly, more than a decade ago, decreased expression of antioxidants and NRF2 levels were described in DRG of the YG8R mouse model [25]. Therefore, to elucidate the potential role of ferroptosis in FA, we analyzed both the ferroptotic traits and the main pathways leading to the NRF2 decline in DRG neurons under frataxin deficiency. For this endeavor, we used two models: primary cultures of frataxin-deficient DRG neurons and DRG isolated from FXN^I151F^ mouse model [26]. This mouse carries the I151F point mutation, equivalent to the human I154F pathological mutation. FXN^I151F^ homozygous mice present very low frataxin levels, biochemical alterations, and neurological deficits (starting at 23 weeks of age) that mimic those observed in patients with FA. Briefly, FXN^I151F^ mice present reduced weight gain from 15 weeks of age onward, and they show decreased motor coordination, forelimb strength, and locomotor activity, as well as gait ataxia [26]. Based on this information, we used FXN^I151F^ mice at two ages: 21 weeks, before neurological symptoms appear, and 39 weeks, when the neurological symptoms are clearly apparent.

In this study, we found iron dysregulation denoted by increased transferrin receptor 1 (TFR1) and heme oxygenase 1 (HMOX1), as well as a decrease in ferritins. In addition, the transporter system Xc (which enters cystine to generate glutathione) and glutathione peroxidase 4 (GPX4) were also decreased, which, in conjunction with the oxidative stress situation, leads to lipid peroxidation. To decipher the main pathways that may explain such alterations, we focused on NRF2, which showed a decrease in its total and nuclear levels. Negative regulators such as Kelch-like ECH-associated protein 1 (KEAP1) and glycogen synthase kinase 3 (GSK3β) were both upregulated, while AMP-activated protein kinase (AMPK), a positive regulator, was impaired. We also analyzed the role of liver kinase 1 (LKB1) as a main upstream kinase of AMPK. In summary, we have comprehensively characterized the molecular alterations caused by frataxin deficiency in DRG neurons and have provided new insights into the regulatory pathways.

## 2. Materials and methods

### 2.1. Animals

All animal experiments were performed according to the National Guidelines for the regulation of the use of experimental laboratory animals issued by the Generalitat de Catalunya and the Government of Spain (article 33.a 214/1997), which comply with the ARRIVE guidelines. The experimental protocols were evaluated and approved by the Experimental Animal Ethical Committee of the University of Lleida (CEEA). All procedures were performed at the animal facility. For euthanasia, the guidelines of the American Veterinary Medical Association (AVMA) were followed. Male and female Sprague Dawley rats (RRID:RGD_70508) were maintained in standard conditions with a 12-h photoperiod, a constant temperature of 20°C, and food and water available *ad libitum*.

FXN^I151F^ heterozygous mice (C57BL/6J-Fxnem10(T146T,I151F)Lutzy/J) were obtained from the Jackson Laboratory (Bar Harbor, ME, USA; Stock Number 31922) as described previously [26]. Intercrosses of heterozygous mice were performed to generate the homozygous wild type (WT) and FXN^I151F^ mice. The animals were housed in standard ventilated cages with a 12-h photoperiod and fed with a normal chow diet available *ad libitum*. The animals were weighed weekly. For genotyping DNA was extracted from tail biopsy specimens and used for polymerase chain reaction (PCR) as described previously [26].

### 2.2. Isolation of DRG from FXN^I151F^ mice

DRG were isolated as described previously (Sanz-Alcázar et al., 2023). Lumbar, thoracic, and cervical DRG were collected, snap-frozen in liquid nitrogen, and stored at -80°C. Approximately 30–40 DRG were obtained per mouse, resulting in an average yield of 200 ± 100 µg of protein.

### 2.3. Isolation and culture of primary rat DRG sensory neurons

DRG were extracted from neonatal Sprague Dawley rats (postnatal day 3 [P3]–P4) and purified as described previously [27] with modifications [18]. After plating cells (10,000–15,000 cells), lentivirus transduction with short hairpin RNA (shRNA) targeting frataxin mRNA was performed, using specific clones (FXN1 and FXN2) and a scrambled sequence control (Scr). Lentivirus particles were added, and experiments were conducted after 3 or 5 days.

### 2.4. Nuclear and cytoplasmic fractionation

Isolated DRG from mice were resuspended in 200 µL of 50 mM Tris-HCl (pH 7.5), 200 μM ethylenediaminetetraacetic acid (EDTA; Sigma), a protease inhibitor cocktail (Roche), and a phosphatase inhibitor (Roche), and homogenized using a Dounce homogenizer. The resulting solution was centrifuged at 800 *g* for 10 min to pellet the nuclei. The supernatant was subjected to a second centrifugation at 8,000 *g* for 15 min; the supernatant was considered to be the cytoplasmic fraction. The nuclear fraction was resuspended in 2% sodium dodecyl sulfate (SDS) in 125 mM Tris-HCl (pH 7.4), with protease and phosphatase inhibitors. Simultaneously, 2% SDS was added to the cytoplasmic fraction. Both fractions were sonicated and heated at 98°C for 5 min. Subsequently, the protein concentration was determined using the BCA assay (Thermo Scientific), and the samples were processed for western blotting.

### 2.5. Western blotting

Primary DRG neurons as well as DRG tissue were homogenized as described previously [18]. Protein extracts (10–15 µg) underwent SDS-polyacrylamide gel electrophoresis, followed by transfer to polyvinylidene difluoride (PVDF) membranes. The membranes were incubated in I-Block (ThermoFisher, T2015) to block nonspecific protein binding. The following primary antibodies were used: TFR1 (1:500, Invitrogen, ref. 13-6800), HMOX1 (1:1,000, Santa Cruz, ref. sc-136960), FTL (1:1,000, Abcam, ref. ab69090), FTH1 (1:1,000, Abcam, ref. ab75973), GPX4 (1:1,000, Santa Cruz, ref. sc-166570), SLC7A11 (1:500, Proteintech, ref. 26864-1-AP), NRF2 (1:1,000, Abcam, ref. ab62352), H3 (1:50,000, Abcam, ref. ab1791), KEAP1 (1:2,000, Proteintech, ref. 10503-2-AP), LKB1 (1:1,000, Santa Cruz, ref. sc-32245), GSK3β pY216 (1:1,000, Abcam, ref. ab75745), GSK3β (1:1,000, Cell Signaling, ref. 1679832S), SIRT1 (1:1,000, Cell Signaling, ref.1679475T), AMPKα pT172 (1:1,000, Cell Signaling, ref. 1672535S), and AMPKα (1:1,000, Cell Signaling, ref.1672532S). After blotting, the membranes were stained with Coomassie brilliant blue (CBB). Total protein staining was quantified with the Image Lab software (Bio-Rad) and used as a loading control for normalization of the target proteins.

### 2.6. Determination or reduced glutathione (GSH) and oxidized glutathione (GSSG)

Primary DRG neurons were seeded in a 96-well white plate (SPL, reference 33696) pre-treated with 0.1 mg/mL of collagen. To quantify the GSH, GSSG, and total GSH levels, the GSH/GSSG-Glo™ Assay (Promega, reference V6612) was performed according to the manufacturer’s instructions. Briefly, on day 5 after lentivirus transduction, cells were washed with cold phosphate-buffered saline (PBS). For each experimental condition (Scr, FXN1, and FXN2), two wells were used: one for total GSH and another for GSSG. A standard curve was constructed to determine the GSH concentration in each sample. The plate was read using the ChemiDoc system to detect chemiluminescence. The data were normalized based on the protein concentration.

### 2.7. Immunofluorescence

A suspension of primary DRG neurons was plated on glass coverslips (Epredia) coated with collagen and placed in a culture dish with neurobasal medium. On day 5 after lentivirus transduction, the cells were fixed with 4% paraformaldehyde (PFA) for 10 min at room temperature. Subsequently, the cells were washed three times in PBS and fixed with cold methanol for 10 min. The cells were washed again and permeabilized with 0.5% Triton X-100 (Sigma) in PBS for 30 min at room temperature. To block nonspecific protein binding, the cells were incubated in 5% bovine serum albumin (BSA; Sigma) and 0.5% Triton in PBS for 30 min. Then, the primary antibody—NRF2 (1:100, Abcam, ref. ab62352)—was diluted in 5% BSA and 0.5% Triton in PBS, added to the cells, and incubated overnight at 4°C. Afterward, the cells were washed three times with PBS and incubated with goat anti-rabbit conjugated to Alexa-Fluor 594 (1:400 Invitrogen, ref. A21428) for 1 h in the dark at room temperature. Nuclei were stained with Hoechst (1:400, Sigma). The cells were washed three times with PBS and mounted with Mowiol (Calbiochem). The cells were imaged using an Olympus FV1000 laser-scanning confocal microscope with a 60× objective and using the Z-stack configuration. Fluorescence intensity was quantified using the ImageJ software (National Institutes of Health, Bethesda, MD, USA).

For DRG tissue processing, mice were anaesthetized by intraperitoneal injection with ketamine/xylazine (100 µL/10 g body weight) and subsequently perfused intra-aortically with 0.9% NaCl followed by 4% PFA in PBS. After perfusion, DRG were dissected and immersed in the same fixative overnight at 4°C. Then, the DRG were cryoprotected in a solution containing 30% sucrose in 100 mM phosphate buffer for 24 h at 4°C. A cryoblock was made of Tissue Freezing Medium (TFM, Electron Microscopy Science, ref. 72592) embedding compound on dry ice. Cryosections with a thickness of 14 μm were obtained using a freezing cryostat (Leica). For immunofluorescence, sections were incubated for 1 h in a permeabilization solution (0.3% Triton in PBS) followed by 1 h with blocking solution (5% BSA and 0.3% Triton in PBS). The sections were incubated with different primary antibodies—glutathione GS-pro (1:50, Santa Cruz, ref. sc-52399), TFR1 (1:50, Invitrogen, ref. 13-6800), or SLC7A11 (1:50, Proteintech, ref. 26864-1-AP)—and incubated overnight at 4°C. After this incubation, the sections were washed tree times with PBS and incubated with secondary antibody for 1 h in the dark at room temperature (1:400 Alexa-Fluor 488 goat anti-mouse, Fisher ref. A11017). Subsequently, the samples were stained with Nissl stain (1:150 NeuroTrace™ 530/615 Red Fluorescent Nissl Stain, Life Technology, ref. N21482). Finally, the samples were washed and mounted with Mowiol (Calbiochem). The sections were examined using a confocal microscope (Olympus FV1000) with a 10× objective and using the Z-stack configuration. Fluorescence intensity was quantified using the ImageJ software.

### 2.8. Oxidative stress analysis: Mitochondrial iron and lipid peroxidation

Mitochondrial iron and lipid peroxidation were determined with the fluorescent probes Mito-Ferro Green (Dojindo, ref.M489-10) and BODIPY™ 581/591 C11 (ThermoFisher, ref. D3861), respectively. Primary DRG neurons were placed in ibiTreat 4-well m-slides (Ibidi, Cat# 80,286) pretreated with 0.1 mg/mL of collagen. Three or four days after lentivirus transduction, the cells were washed with Hank’s buffered saline solution and 10 mM HEPES medium and incubated for 30 min at 37°C with 10 µM BODIPY™ 581/591 C11. Following incubation, the cells were washed and then imaged with a confocal microscope (Olympus FV1000) with a 60× objective and using the Z-stack configuration. All images were acquired with the same laser intensity (with respect to the Scr cells), an image bit-depth of 12 bits, image resolution of 1024 × 1024 pixels, XYZ scan mode, one-way scan direction, and the frame sequential mode. Mitochondrial fluorescence intensity per cellular soma was quantified using the ImageJ software.

DRG were dissected in a Petri dish with GHEBS (137 mM NaCl, 2.6 mM KCl, 25 mM glucose, 25 mM HEPES, and 100 μg/mL penicillin/streptomycin) and carefully cleaned of the nerves. GHEBS plus 1 mg/mL collagenase (Roche) and 1 mg/mL dispase II (Sigma) was added to the DRG, and they were incubated for 60 min at 37°C. Following digestion, the cells were washed and homogenized with Dulbecco’s phosphate-buffered saline (DPBS, Gibco, ref.14190-136) containing 0.5% BSA and 3 mg/mL DNAse I grade II (Roche, Cat# 104159). Once resuspended, the cells were filtered through a 70-µm strainer (pluriSelect). The resulting supernatant was centrifuged at 400 *g* for 5 min, and the pelleted cells were resuspended in DPBS containing 0.5% BSA and Red Blood Cell Lysis Solution (Miltenyi, ref. 130-094-183). Once a homogeneous suspension of cells was achieved, the cells were incubated with 10 µM of Mito-Ferro Green or 10 µM BODIPY 581/591 C11 for 30 min at 37°C in DPBS and 0.5% BSA. Then, the cells were centrifuged, resuspended in DPBS plus 0.5% BSA, and analyzed with flow cytometry. In brief, the cells were run at low pressure on a FACS Canto II (Becton Dickinson) for Mito-Ferro Green staining and on a CytoFlex SRT (Beckman Coulter) for BODIPY staining.

### 2.9. Statistical analysis

All data are presented as the mean ± standard error of the mean. The normality of the data was assessed with the Shapiro–Wilk test. For the data with a normal distribution, Student’s t test was used to compare two groups, whereas multiple groups were compared using one-way analysis of variance (ANOVA) when there was one independent variable or two-way ANOVA when there were two independent variables, followed by Tukey’s multiple-comparisons test. Data that were not normally distributed were compared with the Mann–Whitney U test (for two groups) or the Kruskal–Wallis test followed by Dunn’s multiple comparisons test (for more than two groups). There were no corrections for multiple testing. The representative images were chosen to closely match the average values of the respective groups. A p-value of < 0.05 was considered significant. GraphPad Prism 9.0 (GraphPad Software, Inc., La Jolla, CA, USA) were used for statistical analysis and to prepare graphs.

## 3. Results

### 3.1. Iron dyshomeostasis and lipid peroxidation drive ferroptosis in primary cultures of frataxin-deficient DRG neurons

Published evidence indicates that frataxin deficiency causes an imbalance in iron homeostasis. Iron deposits or accumulation have been observed in frataxin-deficient models [28], but their role in the progression of the disease remains unclear [29]. Thus, although ferroptosis has been recently proposed to be involved in FA [19] [30] [31] [32] there are limited data, and few studies have been performed in DRG, which is the primary site of neurodegeneration.

In this study, we analyzed the role of ferroptosis in FA using two models: (i) primary cultures of DRG neurons with reduced levels of frataxin and, (ii) DRG obtained from FXN^I151F^ mice, which have reduced frataxin levels in all tissues [33]. We cultured primary DRG neurons and transduced them with lentiviral vectors carrying frataxin-interfering shRNA (FXN1 or FXN2) or a non-interfering shRNA (Scr) as a control. On day 5 after transduction, the frataxin level was reduced to 20%–30% in the FXN1 and FXN2 cells, similar to what has been observed in patients with FA [34]. Cell viability also decreased progressively: On day 5, it was 55% and 40% for the FXN1 and FXN2 cells, respectively. Recent studies involving primary cultures of DRG neurons labelled with Mito-Ferro Green showed mitochondrial iron accumulation in frataxin-deficient cells (FXN1 and FXN2) compared with Scr cells [18].

Iron dyshomeostasis in FA is a complex phenomenon. It is generally accepted that frataxin is involved in the synthesis of iron–sulfur clusters as an activator to the cluster assembly inside the mitochondria, and decreased frataxin levels would be sensed by the cells as an iron deficiency. To evaluate this possibility, we analyzed the levels of transferrin receptor 1 (TFR1), the main protein involved in cellular iron uptake (Fig. 1A). The FXN1 and FXN2 cells showed increased levels of TFR1 (1.6- and 2.2-fold, respectively) compared with the Scr cells (Fig. 1B). Ferritin, also important in iron metabolism, is a cytosolic iron-storage protein composed of two subunits, namely ferritin heavy chain 1 (FTH1) and ferritin light chain (FTL). The heavy and light ferritin subunits are assembled into a high-molecular-weight apoferritin shell, which can chelate up to ∼4,500 iron atoms. Ferritin has been described to protect cells against ferroptosis. In this regard, we found decreased levels of both FTH1 (Fig. 1C and D) and FTL (Fig. 1E and F) in the primary cultures of frataxin-deficient DRG neurons (FXN1 and FXN2 cells) compared with the Scr cells. We also analyzed another marker of ferroptosis, namely heme oxygenase 1 (HMOX1). This enzyme catabolizes heme into three products: carbon monoxide (CO), biliverdin, and free iron. HMOX1 levels were increased by 1.5- and 2-fold in the FXN1 and FXN2 cells, respectively (Fig. 1G and H).

**Fig. 1.**
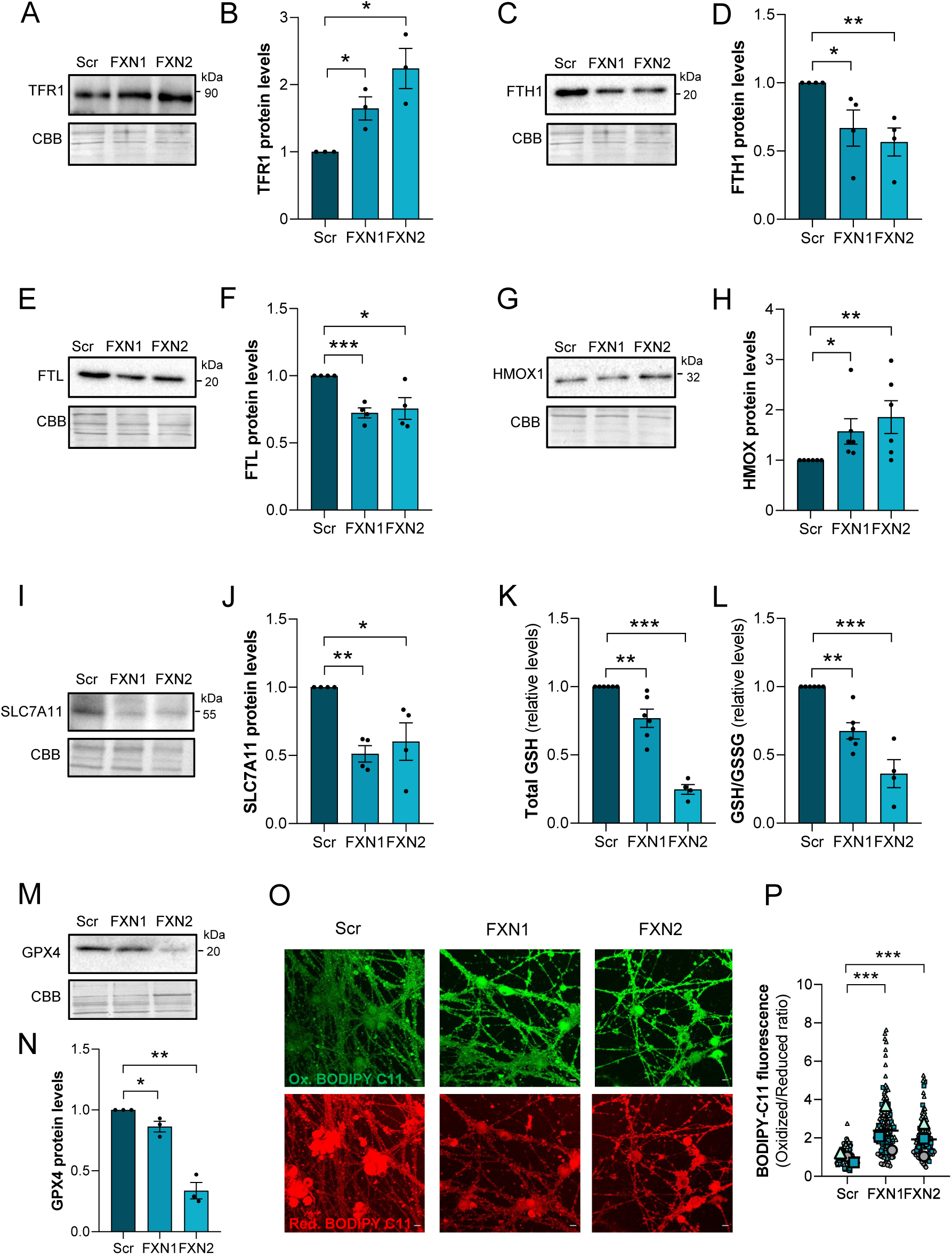
Ferroptotic markers in primary cultures of frataxin-deficient DRG neurons. Scr, FXN1, and FXN2 cells were analyzed at day 5 after lentivirus transduction (unless otherwise stated). Western blotting was used to analyze the indicated proteins. Representative images are shown; CBB protein staining was used as a loading control. (A and B) Western blotting for TRF1; the histograms present the mean ± SEM of n=3 independent cultures. (C and D) Western blotting for FTH1; the histograms present the mean ± SEM of n=4 independent cultures. (E and F) Western blotting for FTL; the histograms present the mean ± SEM of n=4 independent cultures. (G and H) Western blotting for HMOX1; the histograms present the mean ± SEM of n=6 independent cultures. (I and J) Western blotting of SCL7A11; the histograms present the mean ± SEM of n=4 independent cultures. (K and L) Total GSH and the GSH/GSSG ratio were measured; the histograms present the mean ± SEM of n=4–6 independent cultures. (M and N) Western blotting for GPX4; the histograms present the mean ± SEM of n=3 independent cultures. (O) Lipid peroxidation was analyzed with BODIPY C11 at day 4 after lentivirus transduction. Green shows oxidized BODIPY C11 and red shows reduced BODIPY C11. The scale bar is 10 µm. (P) The data are presented as the mean ± SEM of three independent isolations. Between 130 and 160 cells were analyzed for each culture. Each biological replicate is color coded (day to day variability), and small symbols represent individual cells that were analyzed. Significant differences between Scr cells and FXN1 or FXN2 cells are indicated (p values <0.05(*), 0.01(**) or 0.001(***)).

Lipid peroxidation plays a central role in driving ferroptotic cell death and can be caused by impairment of the GSH–GPX4 antioxidant system. GSH is needed for GPX4 activity, which reduces lipid peroxides into lipid alcohols. The synthesis of GSH requires cystine, which enters the cell via system Xc, a transmembrane complex composed of solute carrier family 7 member 11 (SLC7A11) and carrier family 3 member 3 (SLC3A2). Once inside the cell, cystine is reduced to cysteine, where it is mainly used to synthesize GSH. To analyze this pathway, we quantified the levels of SLC7A11 and GSH in primary DRG neurons. As shown in Fig. 1I and J, the cystine carrier SLC7A11 was decreased around 50% in the FXN1 and FXN2 cells compared with the Scr cells. Consistently, the total GSH levels were also decreased in the FXN1 cells and especially in the FXN2 cells, as well as the GSH/GSSG ratio (Fig. 1K and L and Supplementary Fig. S1). In addition, we found a significant decrease in GPX4 in the FXN1 cells and especially in the FXN2 cells compared with the Scr cells (Fig. 1M and N). Decreased GPX4 and GSH levels, in addition to the oxidative stress generated by mitochondrial iron accumulation, would lead to increased lipid peroxidation, a main marker of ferroptosis. We assessed cellular lipid oxidation in primary sensory neurons by using BODIPY C11 581/591. The FXN1 and FXN2 cells showed increased green fluorescence indicative of lipid peroxidation (Fig. 1O), which was quantified as a 2–3-fold increase (Fig. 1P). Taken together, these data clearly indicate that ferroptosis plays a significant role in these frataxin-deficient primary cultures of sensory neurons.

### 3.2. Characterization of ferroptosis in DRG from FXN^I151F^ mice

We wondered whether iron accumulation and ferroptosis also occur in an *in vivo* model of FA. To that end, we isolated DRG from FXN^I151F^ mice (21 and 39 weeks of age) and quantified mitochondrial iron with flow cytometry using Mito-Ferro Green. DRG from FXN^I151F^ mice accumulated more than twice as much mitochondrial as DRG from WT mice at both ages (Fig. 2A). Similarly to primary cultures, TFR1 was also increased in DRG isolated from FXN^I151F^ mice compared with DRG isolated from WT mice, as detected with western blotting (Fig. 2B and C) and immunofluorescence (Fig. 2D and E). In mice, ferritins presented a different behavior. There were decreased levels of FTH1 at 21 and 39 weeks in DRG isolated from FXN^I151F^ mice compared with DRG isolated from WT mice (Fig. 2F and G), similarly to the primary cultures. Interestingly, FTL increased at both ages, although the differences between the FXN^I151F^ and WT mice were more pronounced at 21 weeks (Fig. 2H and I). Like in primary cultures, HMOX1 levels were increased at 21 and 39 weeks in DRG isolated from FXN^I151F^ mice compared with DRG isolated from WT mice (by 1.5- and 1.7-fold, respectively; Fig. 2J and K).

**Fig. 2.**
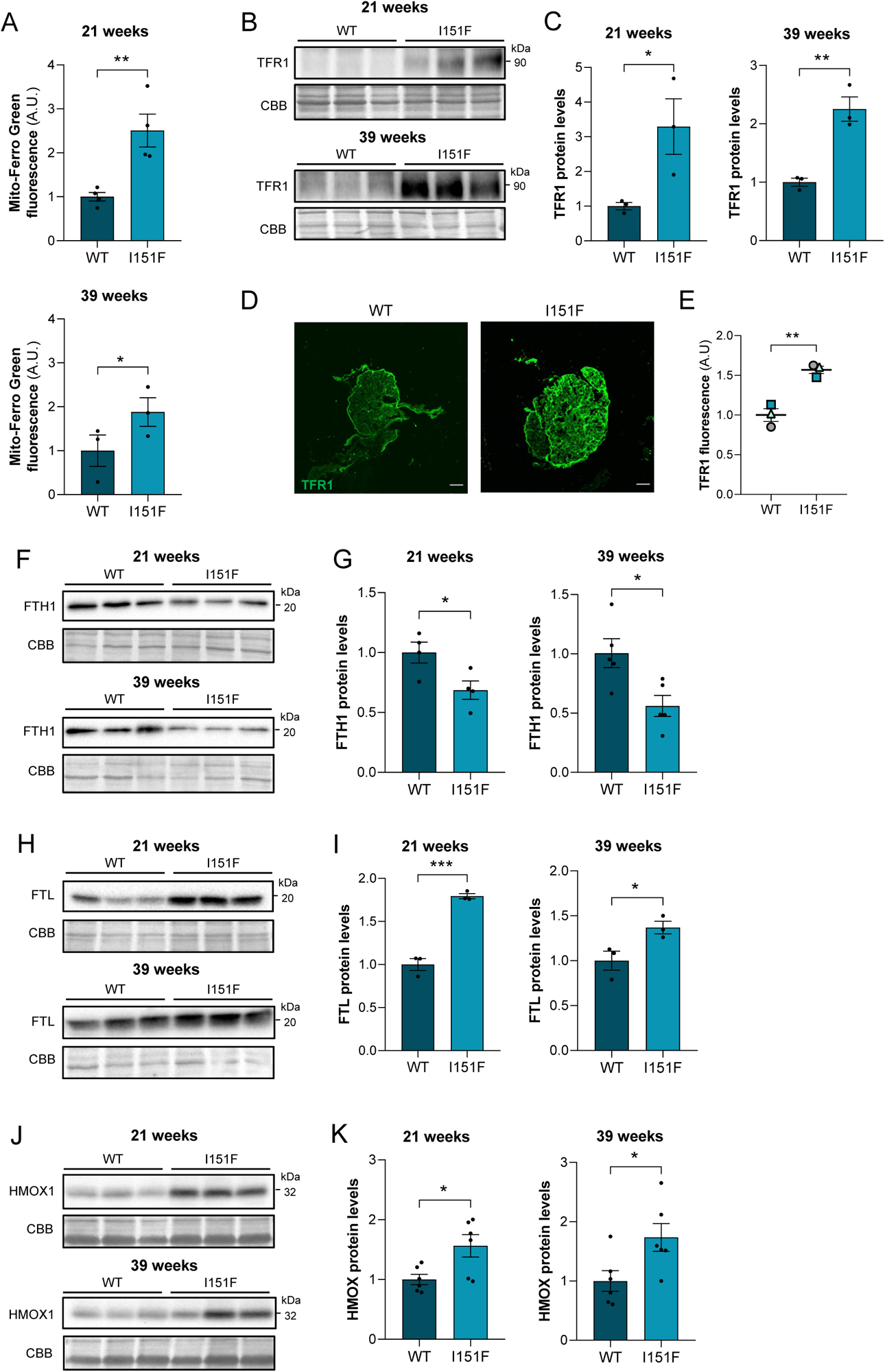
Iron dysregulation in DRG isolated from FXN^I151F^ mice. DRG isolated from 21- and 39-week-old FXN^I151F^ were compared with DRG isolated from WT mice. In each experiment, representative images are shown. (A) Mitochondrial iron (Fe^2+^) was quantified with flow cytometry and the Mito-Ferro Green probe. The data represent the mean ± SEM of n=4 mice per group (21 weeks) or n=3 mice per group (39 weeks). (B) TFR1 protein levels were analyzed by western blotting. (C) The histograms present the mean ± SEM of n=3 mice per group. CBB protein staining was used as a loading control. (D) TFR1 was analyzed by immunofluorescence staining of DRG isolated from 21-week-old mice. The scale bar is 100 μm. (E) The data are presented as the mean ± SEM of n=3 independent isolations. (F) FTH1 protein levels were analyzed by western blotting. (G) The histograms present the mean ± SEM of n=4–5 mice per group. (H) FTL protein levels were analyzed by western blotting. (I) The histograms present the mean ± SEM of n=3 mice per group. (J) HMOX1 protein levels were analyzed by western blotting. (K) The histograms present the mean ± SEM of n=6 mice per group. CBB protein staining was used as a loading control. Significant differences between WT and FXN^I151F^ mice are indicated (p values <0.05(*), 0.01(**) or 0.001(***)).

We analyzed the GSH–GPX4 pathway in FXN^I151F^ mice. As shown in Fig. 3A and B, we found decreased SLC7A11 at 21 weeks in FXN^I151F^ mice compared with WT mice. However, the differences were not significant at 39 weeks. In 21-week-old mice, there was a 50% decrease in SCL7A11 based on immunofluorescence (Fig. 3C and D). GPX4 was reduced at both 21 weeks (by 50%) and 39 weeks (by 25%) in FXN^I151F^ mice compared with WT mice (Fig. 3E and F). We quantified lipid peroxidation with flow cytometry using BODIPY 581/591 C11; it was increased at both ages (Fig. 3G). We also wondered whether, in addition to lipid peroxidation, other markers of oxidative stress could be detected in DRG isolated from FXN^I151F^ mice. Thus, we detected glutathione-protein conjugates by immunofluorescence in DRG isolated from 21-week-old mice, using the GS-Pro antibody. As shown in Fig. 3H and I, protein glutathionylation was increased, by 1.8-fold, in FXN^I151F^ mice compared with WT mice. In summary, the elevated production of hydroperoxide phospholipids in the presence of free iron without proper neutralization by the system Xc/GSH–GPX4 axis indicates a role of ferroptosis in DRG of FXN^I151F^ mice.

**Fig. 3.**
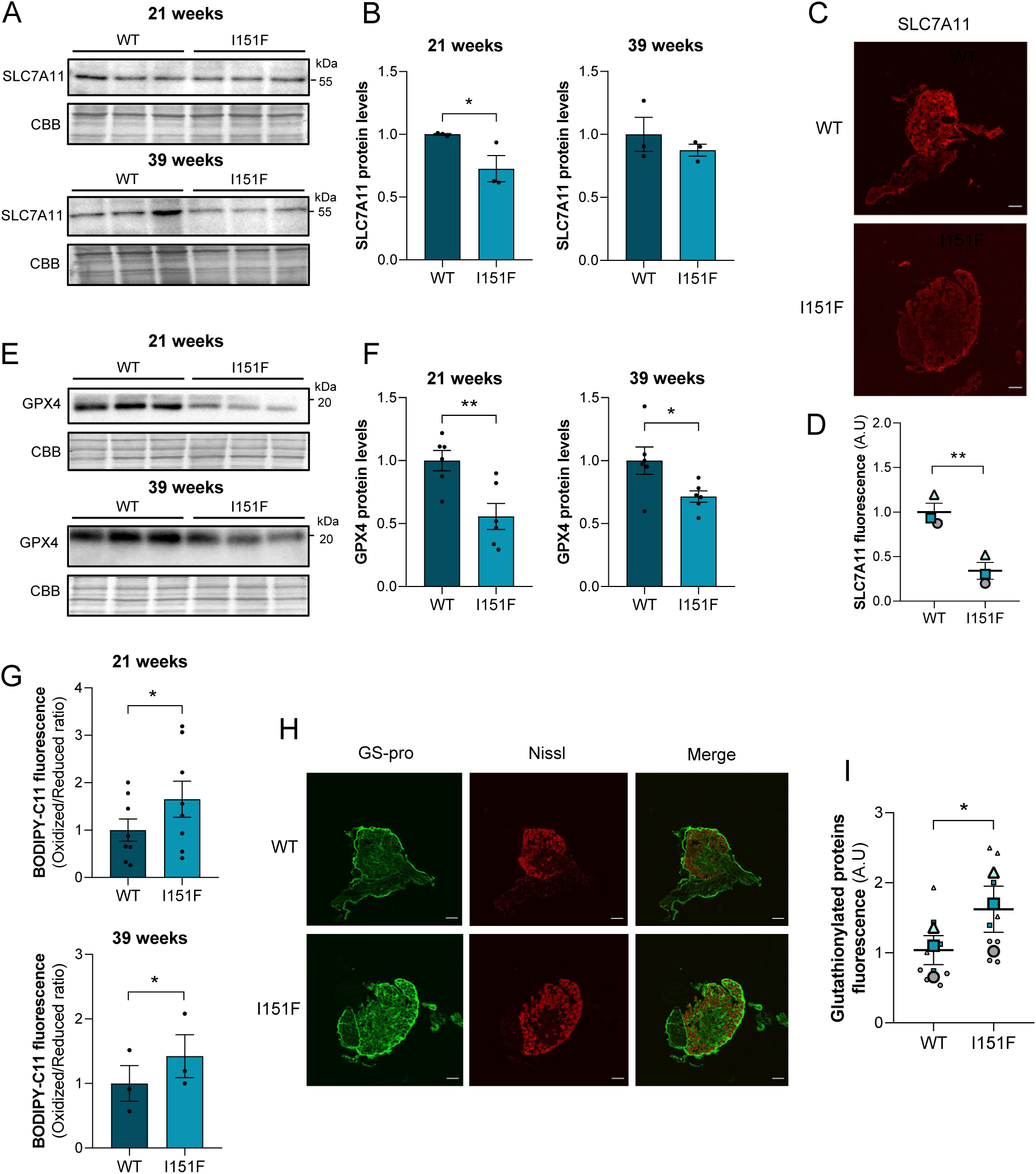
Ferroptotic markers in DRG isolated from FXN^I151F^ mice. DRG isolated from 21- and 39-week-old FXN^I151F^ were compared with DRG isolated from WT mice. In each experiment, representative images are shown. (A) SCL7A11 protein levels were analyzed by western blotting. (B) The histograms present the mean ± SEM of n=3 mice per group. CBB protein staining was used as a loading control. (C) SLC7A11 was analyzed by immunofluorescence staining in DRG isolated from 21-week-old mice. The scale bar is 100 μm. (D) The data are presented as the mean ± SEM of n=3 independent isolations. (E) GPX4 protein levels were analyzed by western blotting. (F) The histograms present the mean ± SEM of n=6 mice per group. CBB protein staining was used as a loading control. (G) Lipid peroxidation was performed by flow cytometry with the BODIPY C11 probe. The ratio of oxidized (510 nm) to reduced (590 nm) BODIPY-C11 is shown. The data are presented as the mean ± SEM of n=6 mice per group. (H) Protein glutathionylation was analyzed by immunohistochemistry in DRG isolated from 21-week-old mice. Glutathionylated proteins (green), Nissl neuronal staining (red), and the merged image are shown. The scale bar is 100 μm. (I) The data are presented as the mean ± SEM fluorescence intensity of the GS-pro antibody of n=3 independent experiments. Each biological replicate is color coded (day to day variability), and small symbols represent different ganglia analyzed in each experiment. Significant differences between WT and FXN^I151F^ mice are indicated (p values <0.05(*), 0.01(**) or 0.001(***)).

### 3.3. Decreased total and nuclear NRF2 in frataxin-deficient DRG neurons: The role of KEAP1 and GSK3β

Although ferroptosis has been associated with FA, the pathways leading to this form of cell death are not fully understood, especially in DRG neurons. A few transcription factors have been described as negative regulators of ferroptosis by controlling the expression of genes involved in iron-related metabolism. Among them, NRF2 is the key player that regulates cytoprotective responses to ferroptotic damage as well as other stresses. We found decreased NRF2 in primary cultures of frataxin-deficient DRG neurons (a 20% and 35% reduction in FXN1 and FXN2 cells, respectively; Fig. 4A and B). Immunofluorescence confirmed the reduction in NRF2 and allowed us to quantify the nuclear and cytosolic localization of this protein. NRF2 decreased in the nucleus and the cytosol (Fig. 4C and D). These results might be partially explained by increased levels of KEAP1, the transcriptional repressor of NRF2. Under physiological conditions, cytosolic NRF2 is bound to KEAP1, allowing NRF2 to be ubiquitinated for proteasomal degradation. In primary cultures, the KEAP1 levels increased by 1.5- and 2.4-fold in FXN1 and FXN2 cells, respectively, compared with Scr cells (Fig. 4E and F). DRG isolated from FXN^I151F^ mice, showed a decrease in total NRF2 at 21 and 39 weeks of age (Fig. 5A and B). In addition, we separated the nuclear and cytosolic fractions (Fig. 5C) and quantified the NRF2 levels in each (Fig. 5D). Nuclear NRF2 was decreased in DRG isolated from FXN^I151F^ mice compared with DRG isolated from WT mice at both ages. However, there were no significant differences in the cytosolic fraction. Although the nuclear/cytosolic NRF2 ratio showed a clear tendency to decrease at both ages, the differences were not significant (Supplementary Fig. S2). Like in primary cultures of DRG neurons, the transcriptional repressor KEAP1 also increased in DRG isolated from FXN^I151F^ mice (1.4- and 1.3-fold at 21 and 39 weeks, respectively) compared with DRG isolated from WT mice (Fig. 5E and F).

**Fig. 4.**
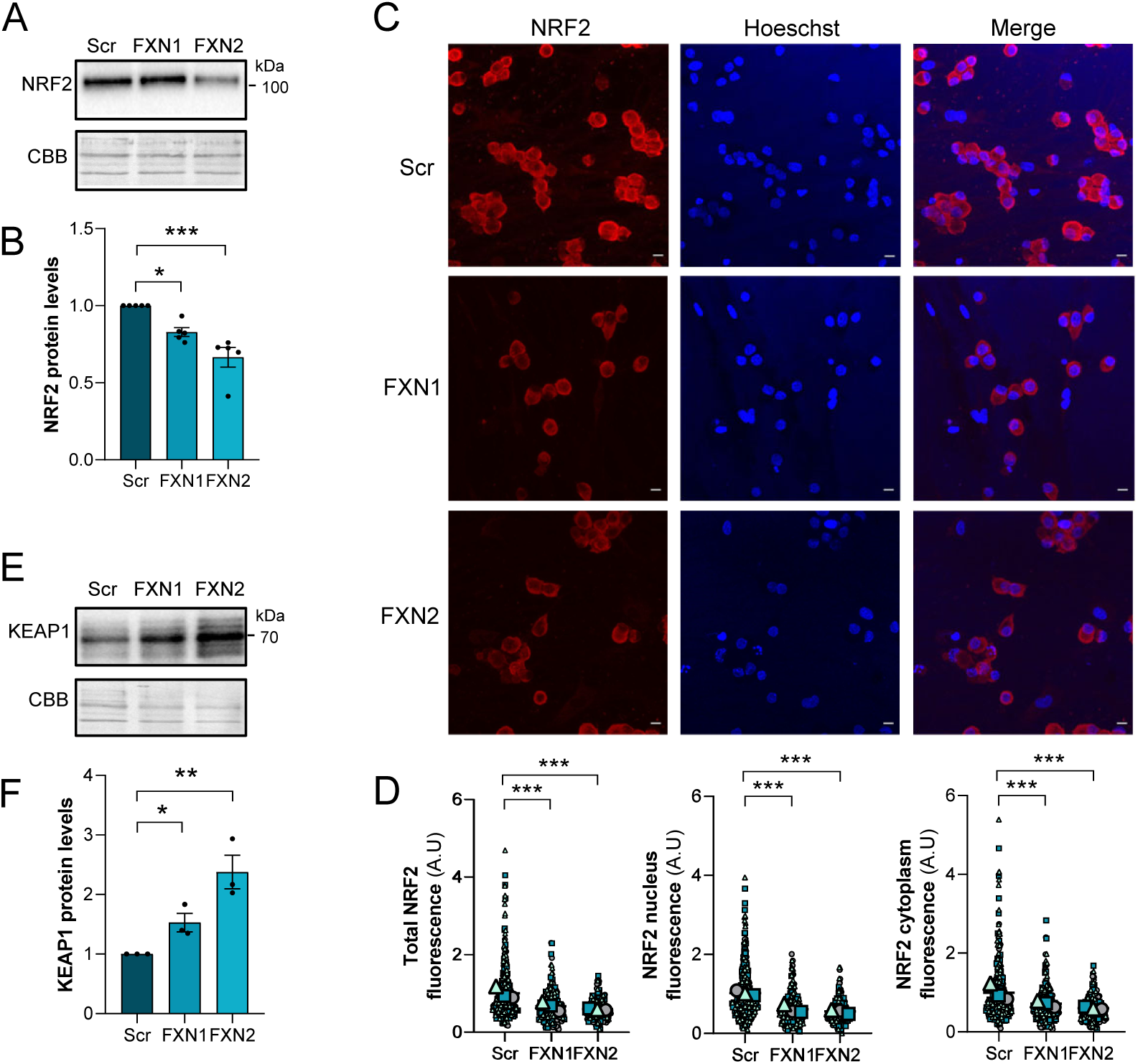
NRF2 and KEAP1 levels in primary cultures of DRG neurons. Frataxin-deficient FXN1 and FXN2 cells were analyzed at day 5 after lentivirus transduction and compared with Scr cells. Western blotting was used to analyze the indicated proteins. Representative images are shown; CBB protein staining was used as a loading control. (A) NRF2 protein levels were analyzed by western blot. (B) The data are presented as the mean ± SEM of n=5 independent cultures. (C) NRF2 was analyzed by immunofluorescence staining. NRF2 (red), Hoechst nuclear staining (blue), and the merged image are shown. The scale bar is 10 µm. (D) The data are presented as the mean ± SEM of n=3 independent experiments. Between 200 and 230 cells were analyzed for each culture. Each biological replicate is color coded (day to day variability), and the small symbols represent different cells that were analyzed. (E) KEAP1 protein levels were analyzed by western blotting. (F) The data are presented as the mean ± SEM of n=3 independent cultures. Significant differences between Scr cells and FXN1 or FXN2 cells are indicated (p values <0.05(*), 0.01(**) or 0.001(***)).

**Fig. 5.**
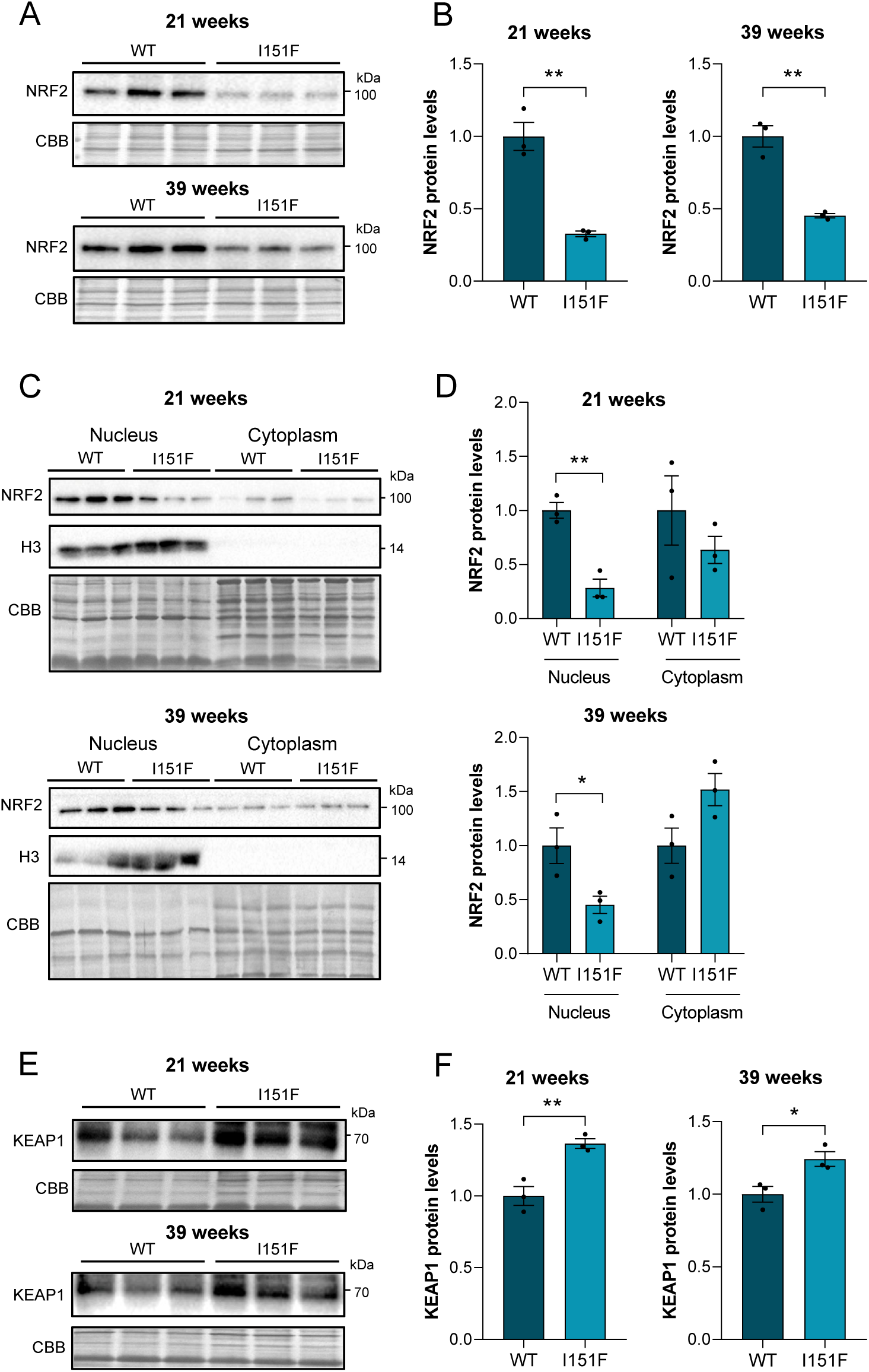
NRF2 and KEAP1 levels in DRG isolated from FXN^I151^ mice. DRG isolated from 21- and 39-week-old FXN^I151F^ were compared with DRG isolated from WT mice. In each experiment, representative images are shown. CBB protein stain was used as a loading control. (A) NRF2 protein levels were analyzed by western blotting using DRG homogenates. (B) The histograms present the mean ± SEM of n=3 mice per group. (C) Nuclear and cytosolic DRG fractions were separated and western blotting was used to detect NRF2. (D) The histograms present the mean ± SEM of n=3 mice per group. (E) KEAP1 protein levels were analyzed by western blotting. (F) The histograms present the mean ± SEM of n=3 mice per group. Significant differences between WT and FXN^I151F^ mice are indicated (p values <0.05(*), 0.01(**) or 0.001(***)).

GSK3β, a multifunctional serine/threonine kinase, is also an NRF2 repressor. Activated GSK3β can phosphorylate NRF2, increasing its nuclear export and degradation [35]. Thus, it could also participate in the decrease in the total and nuclear levels of NRF2 we found. To understand this pathway, we analyzed phosphorylation GSK3β at Tyr216, which leads to activation of this protein, by western blotting in total cell lysates of primary cultures (Fig. 6A) and DRG from mice (Fig. 6C). The pGSK3β/GSK3β ratio was significantly increased in the FXN1 and FXN2 cells (Fig. 6B) and in DRG isolated from FXN^I156F^ mice at 21 and 39 weeks of age (Fig. 6D) compared with their respective controls. These observations indicate that GSK3β is activated (by phosphorylation at Tyr216) and thus decreases NRF2 nuclear localization and promotes its degradation.

**Fig. 6.**
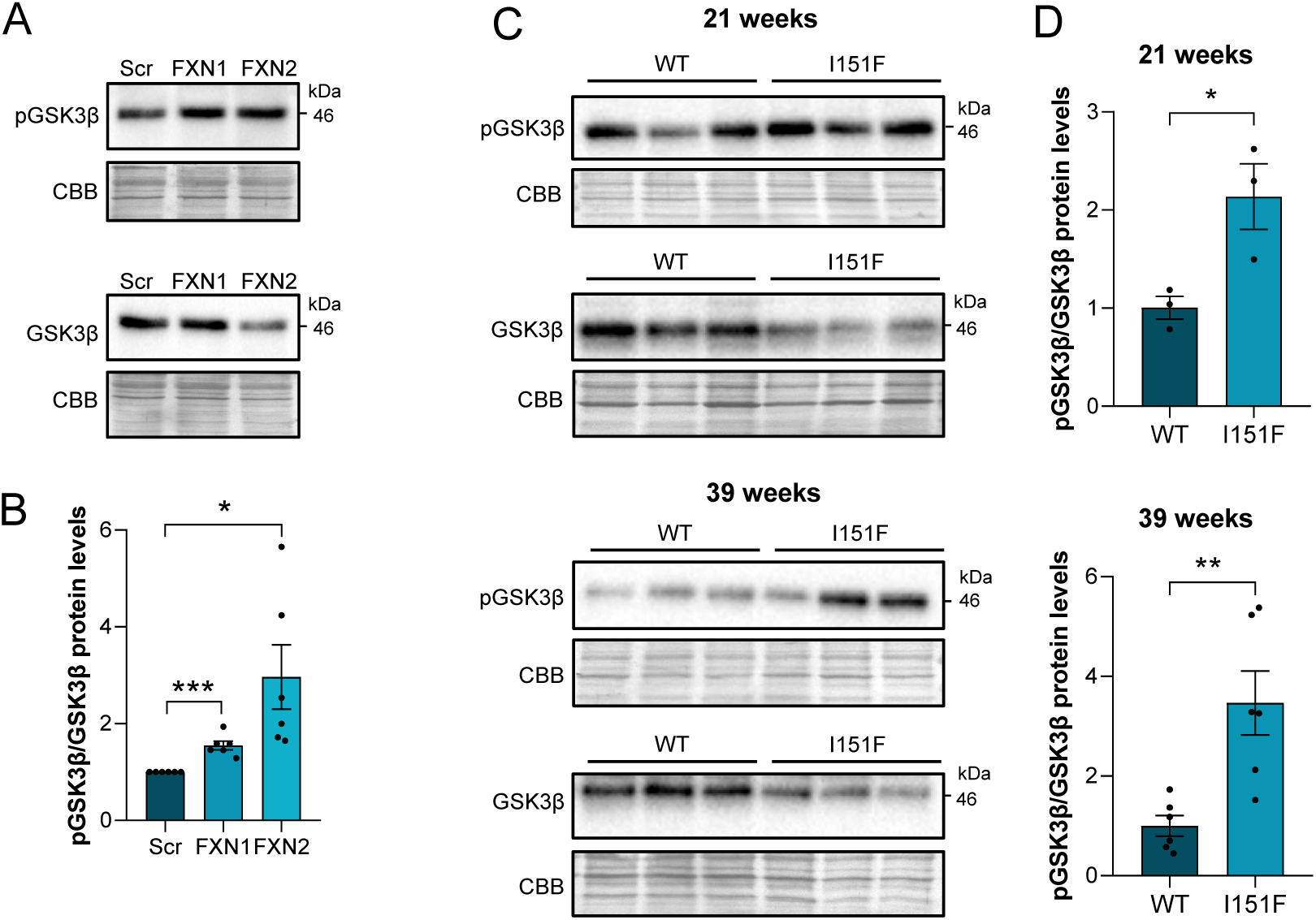
GSK3β in primary cultures of DRG neurons and DRG isolated from FXN^I151F^ mice. (A) GSK3β phosphorylated at Tyr216 and total GSK3β were analyzed by western blotting in Scr (control) as well as FXN1 and FXN2 (5 days after lentivirus transduction) homogenates. Representative western blot images are shown. CBB protein staining was used as a loading control. (B) The histograms present the mean ± SEM of n=6 independent isolations. (C) GSK3β phosphorylated at Tyr216 and total GSK3β were analyzed by western blotting of DRG homogenates from 21- and 39-week-old WT and FXN^I151F^ mice. (D) The histograms represent the mean ± SEM of n=3 mice per group. CBB protein staining was used as a loading control. Significant differences are indicated (p values <0.05(*), 0.01(**) or 0.001(***)).

### 3.4. The LKB1/AMPK signaling cascade in frataxin-deficient DRG neurons

In addition to KEAP1 and GSK3β acting as NRF2 repressors, we wondered whether AMPK could also play a role in NRF2 localization in FA. AMPK, which is considered to be a main cellular energy sensor, facilitates nuclear accumulation of NRF2 though its phosphorylation at Ser550, located in the nuclear export signal (Joo et al., 2016). The mechanism of AMPK activation involves both allosteric modification by AMP and phosphorylation at Thr172 located in the α subunit. AMPK can be phosphorylated by upstream kinases, including LKB1, calcium/calmodulin-dependent protein kinase kinase β (CaMKK2), and transforming growth factor-beta-activated kinase 1 (TAK1) [36].

In primary cultures of DRG neurons, AMPK phosphorylation at Thr172 was decreased (Fig. 7A), with a 40% and 55% reduction in the pAMK/AMPK ratio in FXN1 and FXN2 cells, respectively, compared with Scr cells (Fig. 7B). There were similar results in DRG isolated from FXN^I151F^ mice (Fig. 7C): The pAMPK/AMPK ratio was decreased by 40% at 21 weeks of age and by 60% at 39 weeks of age (Fig. 7D). To investigate the cause of AMPK inactivation, we focused on LKB1, one of the upstream kinases of AMPK. LKB1 is the key component of the mechanisms by which AMPK senses the energy status of the cell [36]. As shown in Fig. 7E and F, the LKB1 levels were markedly reduced by 50% and 80% in FXN1 and FXN2 cells, respectively, compared with Scr cells. However, the LKB1 levels from the total cell lysates did not show significant differences in DRG isolated from 21- or 39-week-old FXN^I151F^ mice compared with DRG isolated from WT mice, although there was a tendency for a decrease at 39 weeks of age (Fig. 7G and H). Given that LKB1 exhibits nucleocytoplasmic shuttling, we evaluated LKB1 in nuclear and cytosolic fractions. As shown in Fig. 7I and J, there was a 40% and 65% decrease in nuclear LKB1 in DRG isolated from 21- and 39-week-old FXN^I151F^ mice, respectively, compared with DRG isolated from WT mice.

**Fig. 7.**
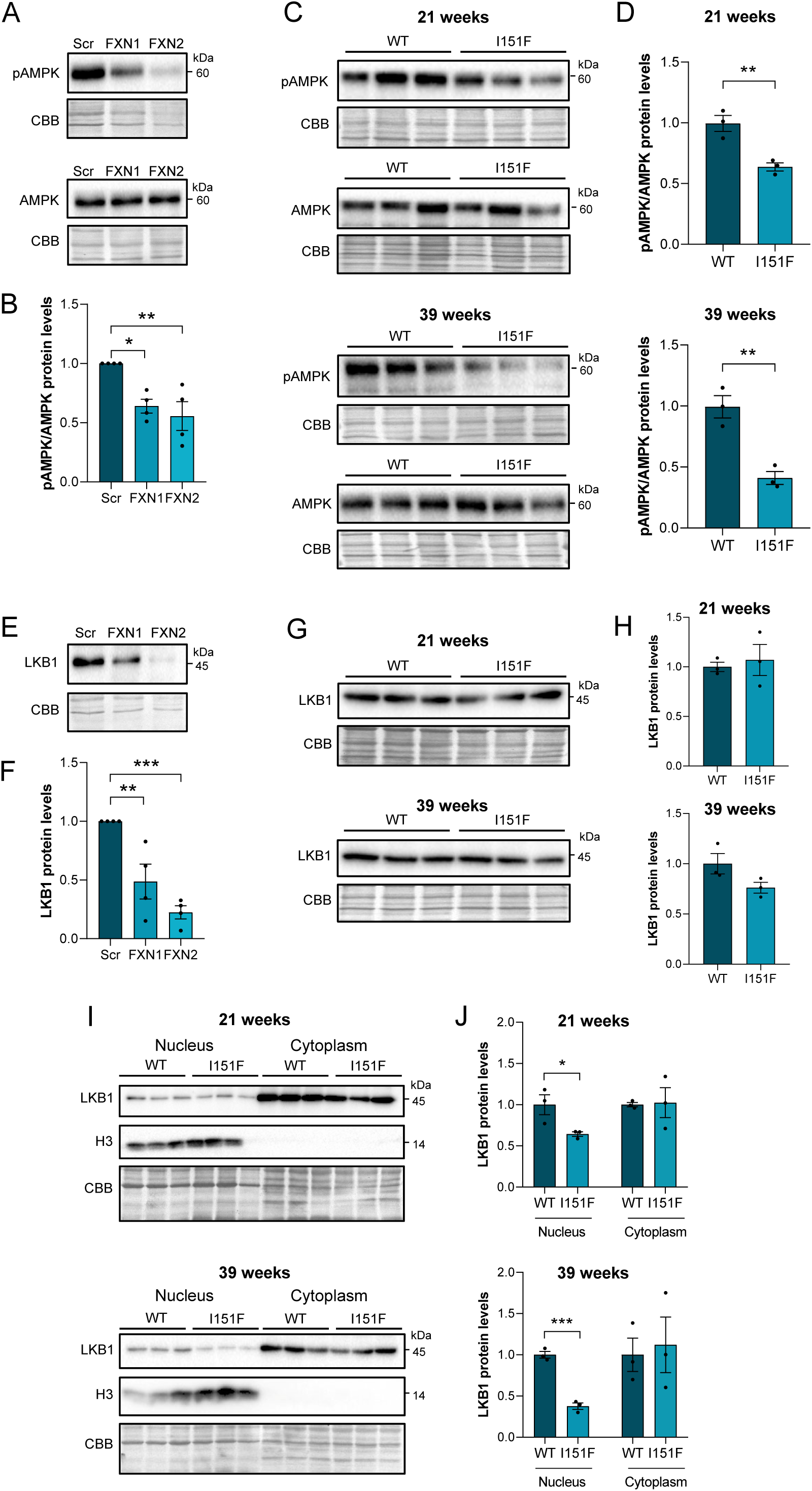
The LKB1/AMPK pathway in primary cultures of DRG neurons and DRG isolated from FXN^I151F^ mice. (A) AMPK phosphorylated at Thr172 and total AMPK were analyzed by western blotting in Scr (control) as well as FXN1 and FXN2 (5 days after lentivirus transduction) homogenates. (B) The histograms present the mean ± SEM of n=4 independent isolations. (C) AMPK phosphorylated at Thr172 and total AMPK were analyzed by western blotting of DRG homogenates from 21- and 39-week-old WT and FXN^I151F^ mice. (D) The histograms present the mean ± SEM of n=3 mice per group. (E) LKB1 was analyzed by western blotting in Scr (control) as well as FXN1 and FXN2 (5 days after lentivirus transduction) homogenates. (F) The histograms present the mean ± SEM of n=4 independent isolations. (G) LKB1 was analyzed by western blotting of DRG homogenates from 21- and 39-week-old WT and FXN^I151F^ mice. (H) The histograms present the mean ± SEM of n=3 mice per group. (I) Nuclear and cytosolic DRG fractions were separated, and LKB1 was analyzed by western blotting. Nuclear histone 3 (H3) was used as a control for fractionation. (J) The histograms present the mean ± SEM from n=3 mice per group. In all figures, representative western blot images are shown. CBB protein staining was used as a loading control. Significant differences are indicated (p values <0.05(*), 0.01(**) or 0.001(***)).

LKB1 is regulated at different levels, including activation via deacetylation by SIRT1 [37]. To study this pathway, we analyzed SIRT1 levels with western blotting of cell lysates from frataxin-deficient primary cultures of DRG neurons (Fig. 8A). There was a 25% and 55% decrease in FXN1 and FXN2 cells, respectively, compared with Scr (Fig. 8B). We also measured SIRT1 levels in DRG isolated from 21- and 39-week-old FXN^I151F^ mice (Fig. 8C). As shown in Fig. 8D, there was a significant decrease in DRG isolated from FXN^I151F^ mice (22% at 21 weeks and 43% at 39 weeks) compared with DRG isolated from WT mice. In addition, our group recently reported decreased SIRT activity in primary cultures of DRG neurons from 21- and 39-week-old FXN^I151F^ mice compared with their controls [18]. Such decreased SIRT1 levels and activity could explain, at least partially, the inactivation of the LKB1/AMPK axis and consequently, the decreased nuclear localization of NRF2.

**Fig. 8.**
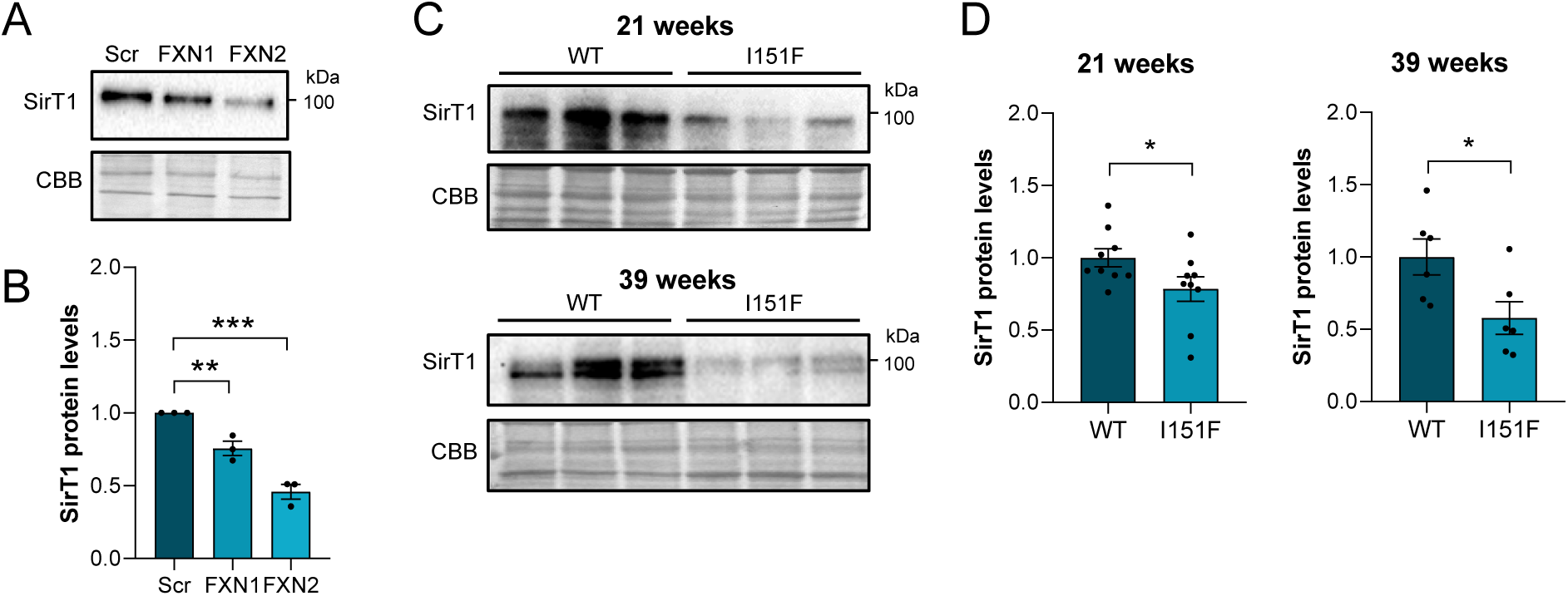
SIRT1 levels in primary cultures of DRG neurons and DRG isolated from FXN^I151F^ mice. (A) SIRT1 was analyzed by western blotting in Scr (control) as well as FXN1 and FXN2 (5 days after lentivirus transduction) homogenates. (B) The histograms present the mean ± SEM of n=3 independent isolations. (C) SIRT1 was analyzed by western blotting of DRG homogenates from 21- and 39-week-old WT and FXN^I151F^ mice. (D) The histograms present the mean ± SEM of n=3 mice per group. In all figures, representative western blot images are shown. CBB protein staining was used as a loading control. Significant differences are indicated (p values <0.05(*), 0.01(**) or 0.001(***)).

## 4. Discussion

It is known that degeneration of DRG large sensory neurons is one of the initial events in FA. However, why DRG are so vulnerable to frataxin deficiency is still unknown. Although iron dysregulation has been described for many years in FA [7] [38], ferroptosis, an iron-dependent form of programmed cell death caused by iron dyshomeostasis and accumulation of lipid peroxides, has recently been reported in cellular models of FA [20] [30] [32] and in heart of the KIKO mouse model [31]. However, no studies have demonstrated whether ferroptosis occurs *in vivo* in DRG sensory neurons. More importantly, the underlying molecular mechanism and regulatory process leading to ferroptosis in FA is not completely known. In the present work, we evaluated the main pathways involved in ferroptosis and the regulatory mechanisms, with a special focus on the transcription factor NRF2, by using two FA models: (i) primary cultures of frataxin-deficient DRG neurons, and (ii) DRG isolated from FXN^I151F^ mice.

In a recent publication, we showed that mitochondrial iron accumulates in frataxin-deficient primary cultures of DRG neurons [18]. Here, we demonstrated mitochondrial iron accumulation *in vivo* in DRG isolated from FXN^I151F^ mice. Such an excess of reduced iron promoted, though the Fenton reaction, oxidative stress. In addition, accumulation of mitochondrial iron is thought to be the result of an increased metabolic need for the metal because, due to frataxin deficiency, its utilization is defective. This may lead to iron flux into the mitochondria, and a cytosolic iron depletion would be sensed by iron-regulatory proteins (IRPs). This phenomenon may also explain why there were no differences in the total iron levels in DRG from patients with FA compared with controls [39]

TFR1-mediated endocytosis is responsible for internalizing iron bound to circulating transferrin. We found increased TFR1 levels in our two models of frataxin-deficient DRG neurons, in agreement with the upregulation of *TFR1* in cardiac tissues from muscle MCK conditional frataxin knockout mice [40]. When cells sense iron starvation, TFR1 is mainly upregulated by association of IRPs to iron-responsive elements (IREs) in the 3′-untranslated region (UTR) of *TFR1* mRNA, increasing its stability [41]. The same system may explain the downregulation of ferritins since they have IREs in the 5′-UTR [42], which leads to decreased translation. Ferritin is a cytosolic iron storage protein composed of two subunits, FTH1 and FTL, assembled into a high-molecular-weight apoferritin shell. FTH1 has ferroxidase activity and can convert ferrous iron (Fe^2+^) to ferric iron (Fe^3+^), which is deposited as ferric hydroxides into the ferritin mineral core that limits iron’s redox switching. Depletion of FTH1 in *Drosophila* larval wing disks leads to ferroptosis-associated severe growth defects, while the depletion of FTL causes only minor defects [43]. These and other findings indicate that ferritin has protective effects on ferroptosis, but the role of FTH1 and FTL on ferroptosis may not be the same [44]. In the present work, decreased ferritin would have a deleterious effect on ferroptosis, but it is still unclear why, in FXN^I151F^ mice, FTL is increased compared with WT mice.

HMOX1 was also upregulated in frataxin-deficient DRG neurons, releasing Fe^2+^ from heme. In most cases, HMOX1 plays a pro-ferroptotic role, and erastin, a ferroptosis inducer, increases the levels of this protein. The HMOX1 inhibitor zinc protoporphyrin IX prevents erastin-induced ferroptotic cell death, whereas the HMOX1 inducer hemin accelerates erastin-induced ferroptosis [45]. HMOX1 knockdown reduced iron overload and ROS, and alleviated lipid peroxidation in diabetic endothelial cells [46]. However, HMOX1 can inhibit ferroptosis in some cases [47], suggesting that the role of HMOX1 in ferroptosis may depend on the context. We observed increased HMOX1 in primary cultures of frataxin-deficient DRG neurons and in DRG isolated from FXN^I151F^ mice. HMOX1 is known to be upregulated by NRF2 [48]; however, we found decreased NRF2 levels in our two models. Such an apparent contradiction has also been observed in cardiac tissues from conditional FXN knockout mice [40], as well as in induced pluripotent stem cell–derived neurons and cardiomyocytes from patients with FA [49]. Nevertheless, it should be kept in mind that HMOX1 can be induced by a variety of stimuli, including heme, cytokines, endotoxins, and heavy metals [44].

Lipid peroxidation plays a key role in ferroptotic cell death, indicative of the collapse of the GSH–GPX4 antioxidant system. The system Xc is a heterodimeric transmembrane complex composed of a light chain, SLC7A11, and a heavy chain, SLC3A2. Cystine enters cells via the system Xc and is subsequently reduced to cysteine, which is mainly used to synthesize GSH. GSH maintains a reduced environment and acts as a potent low-molecular-weight antioxidant inside the cell. It is utilized by GPX4, which uses highly nucleophilic selenocysteine to reduce lipid peroxides into lipid alcohols. In the present study, we found decreased SLC7A11 and GSH levels. The decline in GPX4 would lead to increased lipid peroxidation, one of the main markers of ferroptosis. The reduced SLC7A11 and GPX4 protein levels can be explained by the reduction in nuclear NRF2: The *SLC7A11* and *GPX4* genes have antioxidant-response elements (AREs) in their promoter and are well-known targets of NRF2 [50]. We noted a decreased capacity to reduce lipid oxidation *in vitro* and *in vivo* in the context of frataxin deficiency. In addition, immunohistochemistry revealed increased protein glutathionylation in DRG isolated from FXN^I151F^ mice, which indicates a more oxidized intracellular redox state.

Among the small number of transcription factors that have been reported as negative regulators of ferroptosis, NRF2 plays a key role. Under physiological conditions, NRF2 inhibits the ferroptotic cascade by upregulating genes involved in the maintenance of iron homeostasis and antioxidant defense. Although frataxin deficiency induces oxidative stress inside the mitochondria and the whole cell, defects in NRF2 signaling have been identified in FA fibroblasts [51], motor neurons [52], and in DRG and the cerebella of YG8R hemizygous mouse [25]. In this context, NRF2 activators like omaveloxolone showed to be a promising treatment for FA [53] [54]. In February 2023, Reata Pharmaceuticals announced that the Food and Drug Administration had approved their SKYCLARYS, the brand name for omaveloxolone, for patients with FA aged ≥ 16 years, making it the first approved treatment for FA.

NRF2 exists as a cytoplasmic protein bound to KEAP1; KEAP1 homodimerization and its binding to CULLIN3 allow polyubiquitination of NRF2, promoting its degradation by the ubiquitin proteasomal pathway [55]. When KEAP1 releases NRF2, it translocates to the nucleus and, together with sMAF binding partners, recognize AREs in the promoter of its target genes. In agreement with other FA studies, we found decreased nuclear and cytosolic NRF2 levels in primary cultures of frataxin-deficient DRG neurons. In DRG isolated from FXN^I151F^ mice, there were decreased nuclear NRF2 levels. In both cases, total and nuclear NRF2 levels were reduced, which may explain the decrease in SLC7A11 and GPX4, as noted previously.

Proteins controlling NRF2 nucleocytoplasmic shuttling were altered in our two FA models. An increase in the repressor KEAP1 would be responsible for the decrease in the total NRF2 levels. In the context of FA, increased KEAP1 has been described in fibroblasts of patients [24], as well as in heart of 9-week-old FXN knockout mice [35]. Our results showed, for the first time, an increase in KEAP1 in DRG sensory neurons in *in vitro* and *in vivo* models. This result disagrees with that found in DRG isolated from 6-month-old YG8R hemizygous mouse, namely a reduction in NRF2 levels but no differences in KEAP1 [25]. The differences between our study and the study by Shan et al. (2013) [25] could be explained by the different mouse models and ages. At present, it is unclear why KEAP1 levels are significantly increased, although reduced DJ-1 levels that acts as a stabilizer of NRF2 by preventing the KEAP1–NRF2 association [56], has been reported in fibroblasts from patients with FA [24].

Another negative regulator of NRF2 is GSK3β. In the nucleus, GSK3β is activated by phosphorylation at Tyr216, resulting in direct GSK3β-mediated NRF2 phosphorylation at Ser338 [35]. Such NRF2 phosphorylation induces nuclear accumulation of β-transducin repeat containing E3 ubiquitin protein ligase (β-TRCP), which targets pSer338NRF2 for ubiquitination and subsequent nuclear export and/or degradation. Thus, the increased pTyr216GSK3β/GSK3β ratio we found would lead to phosphorylation, nuclear export, and degradation of NRF2. Moreover, it has been reported that AMPK phosphorylates GSK3β at Ser9, resulting in its inhibition [57]. Because we found decreased AMPK activity (denoted by a decreased pAMPK/AMPK ratio), a decrease in GSK3β phosphorylation at Ser9 is feasible in frataxin-deficient DRG neurons, as has been demonstrated in a mouse model with frataxin knockout in the heart [35]. Taken together, an increase in the pTyr216GSK3/GSK3 ratio and a decrease in AMPK activity would result in GSK3β activation and, consequently, impaired NRF2 signaling.

We found for the first time in FA that the defective NRF2 response may also be mediated by downregulation of the LKB1/AMPK pathway. AMPK is a cellular energetic sensor that, in physiological conditions, is activated by (i) phosphorylation at the Thr172 residue in the α subunit by upstream AMPK kinases and (ii) allosteric modification by AMP when the AMP/ATP ratio is high (indicative of a low energetic state) [58] [59]. Activated AMPK inhibits anabolic pathways, such as lipogenesis, while simultaneously activating catabolic pathways such as fatty acid oxidation, glucose uptake, and mitochondrial biogenesis for efficient ATP generation [60] [61]. Among the main upstream kinases of AMPK, LKB1 encodes a serine/threonine kinase and it is the key component of the mechanisms by which AMPK senses the energy status of the cell. We found reduced levels of LKB1 in DRG sensory neurons both *in vitro* and *in vivo*. Such a decrease can explain the reduced AMPK phosphorylation at Thr172 (and the decrease in the pThr172AMPK/AMPK ratio), negatively regulating AMPK function. AMPK directly phosphorylates NRF2 at Ser550, located in the nuclear export signal, probably inhibiting its nuclear export and leading to NRF2 accumulation in the nucleus [62]. Thus, we hypothesize that in frataxin-deficient DRG neurons, AMPK inactivation reduces NRF2 phosphorylation at Ser550 and, therefore, induces cytoplasmic translocation of NRF2.

LKB1 activity is regulated by deacetylation. Lan et al. [37] demonstrated in cell culture that SIRT1 can deacetylate Lys48 of LKB1 enhancing its kinase activity. In addition, Pillai et al. [63] demonstrated that SIRT3 deacetylates and activates LKB1, thus augmenting the activity of the LKB1/AMPK pathway, and that exogenous NAD^+^ blocks cardiac hypertrophy by activating the SIRT3/LKB1/AMPK pathway. In our previous publication [18], we demonstrated that in DRG sensory neurons, frataxin deficiency leads to electron transport chain impairment; as a consequence, the NAD^+^/NADH ratio, SIRT activity, and SIRT3 levels are decreased. In the present study, we found reduced SIRT1 levels under frataxin deficiency. It is possible that decreased SIRT1 and/or SIRT3 activity and levels increase LKB1 acetylation and inactivation. Consequently, the LKB1/AMPK pathway would be impaired, reducing NRF2 nuclear translocation and activity.

## 5. Conclusion

In conclusion, our study provides further insights regarding how frataxin deficiency induces ferroptosis in sensory neurons as well as in DRG from FXN^I151F^ mice. We observed mitochondrial iron accumulation, accompanied by increased TFR1 and HMOX1 levels as well as decreased ferritin, SLC7A11 and GPX4 levels, leading to oxidative stress and increased lipid peroxidation, all well-known markers of ferroptosis. To elucidate the underlying mechanisms, we investigated the NRF2 signaling pathway, a critical regulator of the cellular response to oxidative stress. We demonstrated increased KEAP1 expression, enhanced activation of GSK3β signaling, and inactivation of the LKB1/AMPK pathway (Fig. 9). Taken together, these results demonstrate why a decrease in the protective activity of NRF2 renders sensory neurons susceptible to ferroptosis.

**Fig. 9.**
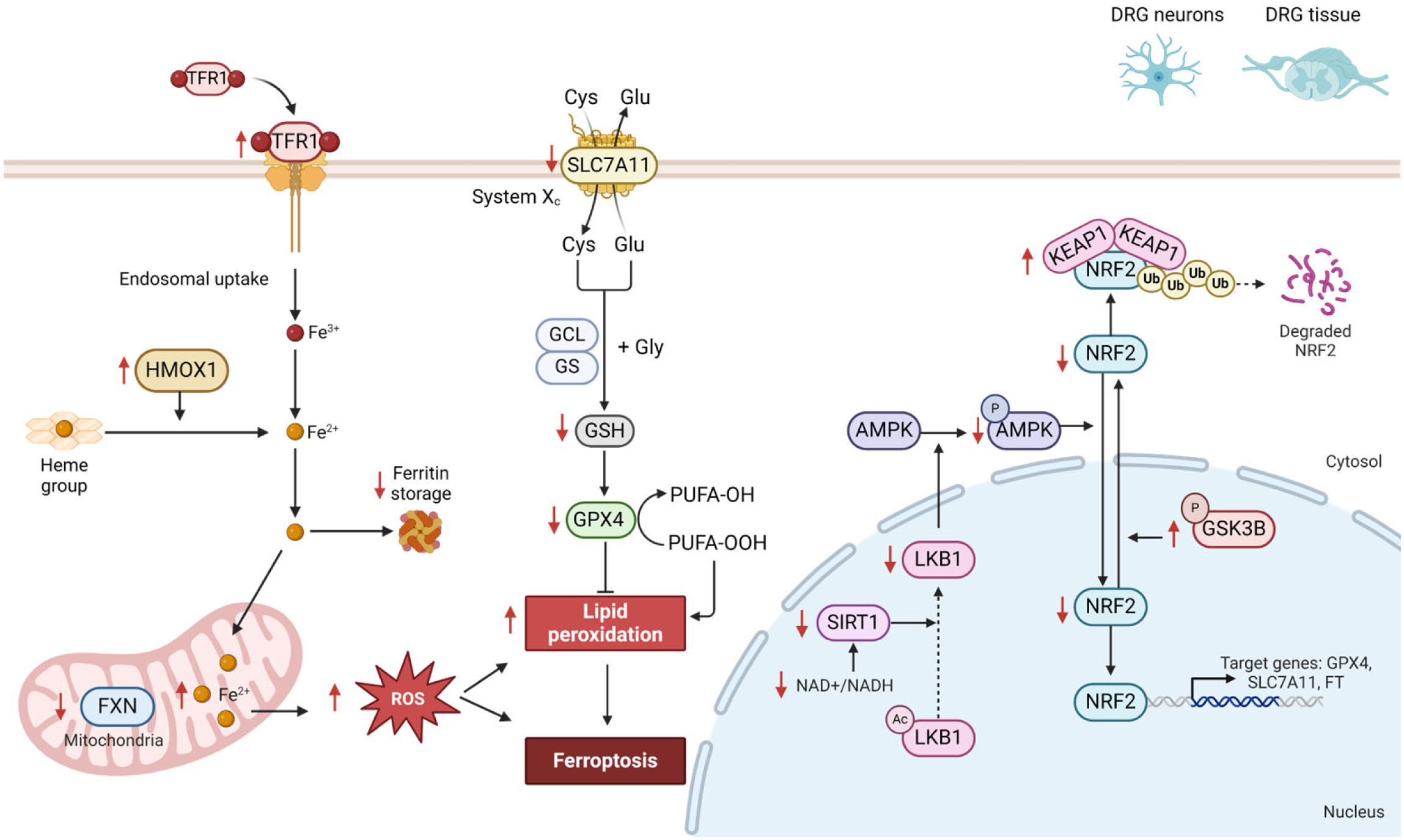
A proposed model of ferroptosis and regulatory pathways in DRG neurons. The figure summarizes the hallmarks of ferroptosis in DRG neurons. The red arrows indicate the results obtained with the two models used in this study. The lack of frataxin causes an accumulation of mitochondrial iron, which is related to an increase in TFR1 and a decrease in FT. This increase in mitochondrial iron contributes to ROS generation and promotes the production of lipid radicals. Increased lipid peroxidation is attributed to reduced levels of SLC7A11, GSH, and GPX4. The decrease in total and nuclear NRF2 can be explained by two parallel processes: (i) increased KEAP1 and pGSK3β, which are involved in the degradation of NRF2, and (ii) decreased pAMPK, LKB1, and SIRT1, which are involved in the nuclear translocation and activation of NRF2.

## Ethics statement

Animal use and care protocol conforms to the National Guidelines for the regulation of the use of experimental laboratory animals from the Generalitat de Catalunya and the Government of Spain (article 33.a 214/1997) and was evaluated and approved by the Experimental Animal Ethical Committee of the University of Lleida (CEEA).

## Acknowledgments

We thank Roser Pané for her excellent technical assistance and Anaïs Panosa, from the Microscopy and Flow Cytometry facilities, for helping in confocal imaging and flow cytometry analysis. We are also grateful to Ester Vilapriñó for her assistance in the statistical analyses.

## Author contributions

Conceptualization, design and perform experiments: A.S-A., F.D., J.T., M.M-C, M.P-G, J.R. and E.C. Data analysis: A.S-A, JR, and E.C. Primary cell cultures and mice control: A.S-A, M.P-C., M.P-G and F.D. Writing original draft: A.S-A, and E.C. Reviewed and editing: J.T., J.R. and E.C. All authors have read and agreed to the published version of the manuscript.

## Data and materials availability

All data required to evaluate the conclusions in the paper are present in the manuscript and the supplementary materials.

## Abbreviations

AMPK: AMP-activated protein kinase
DRG: Dorsal root ganglia
FA: Friedreich ataxia
FTH1: ferritin heavy chain
FTL: ferritin light chain
GPX4: glutathione peroxidase 4
GSH: glutathione
GSK3β: glycogen synthase kinase 3β
HMOX1: heme oxygenase 1
KEAP1: Kelch-like ECH-associated protein 1
LKB1: liver kinase 1
NRF2: nuclear factor erythroid 2–related factor 2
ROS: reactive oxygen species
SOD2: superoxide dismutase 2
TFR1: transferrin receptor 1

## References

[1] V. Campuzano, L. Montermini, M.D. Molto, L. Pianese, M. Cossee, F. Cavalcanti, E. Monros, F. Duclos, A. Monticelli, F. Zara, J. Cafiizares, H. Koutnikova, S.I. Bidichandani, C. Gellera, A. Brice, P. Trouillas, G. De Michele, A. Filla, R. De Frutos, P.I. Patel, S. Di Donato, J. Mandel, S. Cocozza, M. Koenig, M. Pandolfol, Friedreich’s Ataxia: Autosomal Recessive Disease Caused by an lntronic GAA Triplet Repeat Expansion, Science (1979) 271 (1996) 1423–1427.

[2] C.A. Galea, A. Huq, P.J. Lockhart, G. Tai, L.A. Corben, E.M. Yiu, L.C. Gurrin, D.R. Lynch, S. Gelbard, A. Durr, F. Pousset, M. Parkinson, R. Labrum, P. Giunti, S.L. Perlman, M.B. Delatycki, M. V. Evans-Galea, Compound heterozygous FXN mutations and clinical outcome in friedreich ataxia, Ann Neurol 79 (2016) 485–495. 10.1002/ana.24595.

[3] M.H. Parkinson, S. Boesch, W. Nachbauer, C. Mariotti, P. Giunti, Clinical features of Friedreich’s ataxia: classical and atypical phenotypes, J Neurochem 126 (2013) 103–117. 10.1111/JNC.12317.

[4] A.Y. Tsou, E.K. Paulsen, S.J. Lagedrost, S.L. Perlman, K.D. Mathews, G.R. Wilmot, B. Ravina, A.H. Koeppen, D.R. Lynch, Mortality in Friedreich Ataxia, J Neurol Sci 307 (2011) 46–49. 10.1016/J.JNS.2011.05.023.

[5] A. Dürr, M. Cossee, Y. Agid, V. Campuzano, C. Mignard, C. Penet, J.-L. Mandel, A. Brice, M. Koenig, Clinical and genetic abnormalities in patients with Friedreich’s ataxia, N Engl J Med 335 (1996) 1169–1175. 10.1056/NEJM199610173351601.

[6] D. Alsina, R. Purroy, J. Ros, J. Tamarit, Iron in friedreich ataxia: A central role in the pathophysiology or an epiphenomenon?, Pharmaceuticals 11 (2018). 10.3390/ph11030089.

[7] A.H. Koeppen, Tissue iron in Friedreich Ataxia, Journal of Integrative Neuroscience 2024, 23(1), 4 23 (2024) 4. 10.31083/J.JIN2301004.

[8] J. Tamarit, E. Britti, F. Delaspre, M. Medina-Carbonero, A. Sanz-Alcázar, E. Cabiscol, J. Ros, Mitochondrial iron and calcium homeostasis in Friedreich ataxia, IUBMB Life 73 (2021) 543–553. 10.1002/IUB.2457.

[9] A. Moreno-Cermeño, D. Alsina, E. Cabiscol, J. Tamarit, J. Ros, Metabolic remodeling in frataxin-deficient yeast is mediated by Cth2 and Adr1, Biochim Biophys Acta Mol Cell Res 1833 (2013) 3326–3337. 10.1016/j.bbamcr.2013.09.019.

[10] M. Pandolfo, Molecular genetics and pathogenesis of Friedreich ataxia, Neuromuscul Disord 8 (1998) 409–415. 10.1016/S0960-8966(98)00039-X.

[11] D. Simon, H. Seznec, A. Gansmuller, N. Carelle, P. Weber, D. Metzger, P. Rustin, M. Koenig, H. Puccio, Friedreich ataxia mouse models with progressive cerebellar and sensory ataxia reveal autophagic neurodegeneration in dorsal root ganglia, J Neurosci 24 (2004) 1987–1995. 10.1523/JNEUROSCI.4549-03.2004.

[12] M.Z. McMackin, B. Durbin-Johnson, M. Napierala, J.S. Napierala, L. Ruiz, E. Napoli, S. Perlman, C. Giulivi, G.A. Cortopassi, Potential biomarker identification for Friedreich’s ataxia using overlapping gene expression patterns in patient cells and mouse dorsal root ganglion, PLoS One 14 (2019). 10.1371/JOURNAL.PONE.0223209.

[13] A.H. Koeppen, Friedreich’s ataxia: pathology, pathogenesis, and molecular genetics, J Neurol Sci 303 (2011) 1–12. 10.1016/J.JNS.2011.01.010.

[14] A.H. Koeppen, A.B. Becker, J. Qian, B.B. Gelman, J.E. Mazurkiewicz, Friedreich Ataxia: developmental failure of the dorsal root entry zone, J Neuropathol Exp Neurol 76 (2017) 969–977. 10.1093/JNEN/NLX087.

[15] E. Britti, F. Delaspre, J. Tamarit, J. Ros, Calpain-inhibitors protect frataxin-deficient dorsal root ganglia neurons from loss of mitochondrial Na+/Ca2+ exchanger, NCLX, and apoptosis, Neurochem Res 46 (2021) 108–119. 10.1007/s11064-020-03020-3.

[16] R. Purroy, E. Britti, F. Delaspre, J. Tamarit, J. Ros, Mitochondrial pore opening and loss of Ca2 + exchanger NCLX levels occur after frataxin depletion, Biochim Biophys Acta Mol Basis Dis 1864 (2018) 618–631. 10.1016/j.bbadis.2017.12.005.

[17] S. Mincheva-Tasheva, E. Obis, J. Tamarit, J. Ros, Apoptotic cell death and altered calcium homeostasis caused by frataxin depletion in dorsal root ganglia neurons can be prevented by BH4 domain of Bcl-x L protein, Hum Mol Genet 23 (2014) 1829–1841. 10.1093/hmg/ddt576.

[18] A. Sanz-Alcázar, E. Britti, F. Delaspre, M. Medina-Carbonero, M. Pazos-Gil, J. Tamarit, J. Ros, E. Cabiscol, Mitochondrial impairment, decreased sirtuin activity and protein acetylation in dorsal root ganglia in Friedreich Ataxia models, Cell Mol Life Sci 81 (2023) 12. 10.1007/S00018-023-05064-4.

[19] G.M. Cotticelli, S. Xia, D. Lin, T. Lee, L. Terrab, P. Wipf, D.M. Huryn, R.B. Wilson, Ferroptosis as a novel therapeutic target for Friedreich’s Ataxia, J Pharmacol Exp Ther 369 (2019) 47–54. 10.1124/JPET.118.252759.

[20] J. Du, Y. Zhou, Y. Li, J. Xia, Y. Chen, S. Chen, X. Wang, W. Sun, T. Wang, X. Ren, X. Wang, Y. An, K. Lu, W. Hu, S. Huang, J. Li, X. Tong, Y. Wang, Identification of Frataxin as a regulator of ferroptosis, Redox Biol 32 (2020). 10.1016/j.redox.2020.101483.

[21] J. Li, F. Cao, H. liang Yin, Z. jian Huang, Z.T. Lin, N. Mao, B. Sun, G. Wang, Ferroptosis: past, present and future, Cell Death Dis 11 (2020). 10.1038/S41419-020-2298-2.

[22] S.J. Dixon, K.M. Lemberg, M.R. Lamprecht, R. Skouta, E.M. Zaitsev, C.E. Gleason, D.N. Patel, A.J. Bauer, A.M. Cantley, W.S. Yang, B. Morrison, B.R. Stockwell, Ferroptosis: an iron-dependent form of nonapoptotic cell death, Cell 149 (2012) 1060–1072. 10.1016/J.CELL.2012.03.042.

[23] S. Doll, M. Conrad, Iron and ferroptosis: A still ill-defined liaison, IUBMB Life 69 (2017) 423–434. 10.1002/IUB.1616.

[24] S. Petrillo, J. D’amico, P. La Rosa, E.S. Bertini, F. Piemonte, Targeting NRF2 for the treatment of Friedreich’s Ataxia: a comparison among drugs, Int J Mol Sci 20 (2019). 10.3390/IJMS20205211.

[25] Y. Shan, R.A. Schoenfeld, G. Hayashi, E. Napoli, T. Akiyama, M. Iodi-Carstens, E.E. Carstens, M.A. Pook, G.A. Cortopassi, Frataxin deficiency leads to defects in expression of antioxidants and Nrf2 expression in dorsal root ganglia of the Friedreich’s Ataxia YG8R mouse model, Antioxid Redox Signal 19 (2013) 1481– 1493. 10.1089/ars.2012.4537.

[26] M. Medina-Carbonero, A. Sanz-Alcázar, E. Britti, F. Delaspre, E. Cabiscol, J. Ros, J. Tamarit, Mice harboring the FXN I151F pathological point mutation present decreased frataxin levels, a Friedreich ataxia-like phenotype, and mitochondrial alterations, Cellular and Molecular Life Sciences 79 (2022) 1–20. 10.1007/s00018-021-04100-5.

[27] E. Britti, F. Delaspre, A. Feldman, M. Osborne, H. Greif, J. Tamarit, J. Ros, Frataxin-deficient neurons and mice models of Friedreich ataxia are improved by TAT-MTScs-FXN treatment, Journal of Cellular Molecular Medicine 22 (2018) 834–848. 10.1111/jcmm.13365.

[28] J. Tamarit, E. Britti, F. Delaspre, M. Medina-Carbonero, A. Sanz-Alcázar, E. Cabiscol, J. Ros, Crosstalk between nucleus and mitochondria in human disease: Mitochondrial iron and calcium homeostasis in Friedreich ataxia, in: IUBMB Life, Blackwell Publishing Ltd, 2021: pp. 543–553. 10.1002/iub.2457.

[29] J.V. Llorens, S. Soriano, P. Calap-Quintana, P. Gonzalez-Cabo, M.D. Moltó, The role of iron in Friedreich’s Ataxia: insights from studies in human tissues and cellular and animal models, Front Neurosci 13 (2019). 10.3389/FNINS.2019.00075.

[30] P. La Rosa, S. Petrillo, M.T. Fiorenza, E.S. Bertini, F. Piemonte, Ferroptosis in Friedreich’s Ataxia: a metal-induced neurodegenerative disease, Biomolecules 10 (2020) 1–15. 10.3390/BIOM10111551.

[31] P. La Rosa, S. Petrillo, R. Turchi, F. Berardinelli, T. Schirinzi, G. Vasco, D. Lettieri-Barbato, M.T. Fiorenza, E.S. Bertini, K. Aquilano, F. Piemonte, The Nrf2 induction prevents ferroptosis in Friedreich’s Ataxia, Redox Biol 38 (2021). 10.1016/J.REDOX.2020.101791.

[32] R. Turchi, R. Faraonio, D. Lettieri-Barbato, K. Aquilano, An overview of the ferroptosis hallmarks in Friedreich’s Ataxia, Biomolecules 10 (2020) 1489–1504. 10.3390/biom10111489.

[33] M. Medina-Carbonero, A. Sanz-Alcázar, E. Britti, F. Delaspre, E. Cabiscol, J. Ros, J. Tamarit, Biochemical alterations precede neurobehavioral deficits in a novel mouse model of Friedreich ataxia, BioRxiv (2021). 10.1101/2021.04.05.438486.

[34] E. Britti, F. Delaspre, A. Sanz-Alcázar, M. Medina-Carbonero, M. Llovera, R. Purroy, S. Mincheva-Tasheva, J. Tamarit, J. Ros, Calcitriol increases frataxin levels and restores mitochondrial function in cell models of Friedreich Ataxia, Biochem J 478 (2021) 1–20. 10.1042/BCJ20200331.

[35] A. Anzovino, S. Chiang, B.E. Brown, C.L. Hawkins, D.R. Richardson, M.L.H. Huang, Molecular alterations in a mouse cardiac model of Friedreich Ataxia: an impaired Nrf2 response mediated via upregulation of Keap1 and activation of the Gsk3β axis, Am J Pathol 187 (2017) 2858–2875. 10.1016/j.ajpath.2017.08.021.

[36] A. Woods, S.R. Johnstone, K. Dickerson, F.C. Leiper, L.G.D. Fryer, D. Neumann, U. Schlattner, T. Wallimann, M. Carlson, D. Carling, LKB1 is the upstream kinase in the AMP-activated protein kinase cascade, Curr Biol 13 (2003) 2004–2008. 10.1016/J.CUB.2003.10.031.

[37] F. Lan, J.M. Cacicedo, N. Ruderman, Y. Ido, SIRT1 Modulation of the acetylation status, cytosolic localization, and activity of LKB1. Possible role in AMP-activated protein kinase activation, Journal of Biological Chemistry 283 (2008) 27628– 27635. 10.1074/jbc.M805711200.

[38] A.H. Koeppen, E.C. Kuntzsch, S.T. Bjork, R.L. Ramirez, J.E. Mazurkiewicz, P.J. Feustel, Friedreich ataxia: Metal dysmetabolism in dorsal root ganglia, Acta Neuropathol Commun 2 (2014) 1–10. 10.1186/2051-5960-1-26/FIGURES/7.

[39] A.H. Koeppen, J.E. Mazurkiewicz, Friedreich ataxia: neuropathology revised., J Neuropathol Exp Neurol 72 (2013) 78–90. 10.1097/NEN.0b013e31827e5762.

[40] M.L.H. Huang, E.M. Becker, M. Whitnall, Y.S. Rahmanto, P. Ponka, D.R. Richardson, Elucidation of the mechanism of mitochondrial iron loading in Friedreich’s ataxia by analysis of a mouse mutant, Proc Natl Acad Sci U S A 106 (2009) 16381–16386. 10.1073/PNAS.0906784106.

[41] M. Miyazawa, A.R. Bogdan, K. Hashimoto, Y. Tsuji, Regulation of transferrin receptor-1 mRNA by the interplay between IRE-binding proteins and miR-7/miR-141 in the 3’-IRE stem-loops, RNA 24 (2018) 468–479. 10.1261/RNA.063941.117.

[42] N. Wilkinson, K. Pantopoulos, The IRP/IRE system in vivo: insights from mouse models, Front Pharmacol 5 (2014) 176. 10.3389/FPHAR.2014.00176.

[43] S. Mumbauer, J. Pascual, I. Kolotuev, F. Hamaratoglu, Ferritin heavy chain protects the developing wing from reactive oxygen species and ferroptosis, PLoS Genet 15 (2019). 10.1371/JOURNAL.PGEN.1008396.

[44] X. Chen, C. Yu, R. Kang, D. Tang, Iron metabolism in ferroptosis, Front Cell Dev Biol 8 (2020) 590226. 10.3389/fcell.2020.590226.

[45] M.Y. Kwon, E. Park, S.J. Lee, S.W. Chung, Heme oxygenase-1 accelerates erastin-induced ferroptotic cell death, Oncotarget 6 (2015) 24393–24403. 10.18632/ONCOTARGET.5162.

[46] Z. Meng, H. Liang, J. Zhao, J. Gao, C. Liu, X. Ma, J. Liu, B. Liang, X. Jiao, J. Cao, Y. Wang, HMOX1 upregulation promotes ferroptosis in diabetic atherosclerosis, Life Sci 284 (2021). 10.1016/J.LFS.2021.119935.

[47] O. Adedoyin, R. Boddu, A. Traylor, J.M. Lever, S. Bolisetty, J.F. George, A. Agarwal, Heme oxygenase-1 mitigates ferroptosis in renal proximal tubule cells, Am J Physiol Renal Physiol 314 (2018) F702–F714. 10.1152/ajprenal.00044.2017.

[48] M. Jaganjac, L. Milkovic, S.B. Sunjic, N. Zarkovic, The NRF2, thioredoxin, and glutathione system in tumorigenesis and anticancer therapies., Antioxidants (Basel) 9 (2020) 1–41. 10.3390/antiox9111151.

[49] M.B. Angulo, A. Bertalovitz, M.A. Argenziano, J. Yang, A. Patel, T. Zesiewicz, T. V. McDonald, Frataxin deficiency alters gene expression in Friedreich ataxia derived IPSC-neurons and cardiomyocytes, Mol Genet Genomic Med 11 (2023). 10.1002/MGG3.2093.

[50] F. He, X. Ru, T. Wen, NRF2, a transcription factor for stress response and beyond, Int J Mol Sci 21 (2020) 1–23. 10.3390/IJMS21134777.

[51] V. Paupe, E.P. Dassa, S. Goncalves, F. Auchère, M. Lönn, A. Holmgren, P. Rustin, Impaired nuclear Nrf2 translocation undermines the oxidative stress response in Friedreich ataxia, PLoS One 4 (2009). 10.1371/JOURNAL.PONE.0004253.

[52] V. D’Oria, S. Petrini, L. Travaglini, C. Priori, E. Piermarini, S. Petrillo, B. Carletti, E. Bertini, F. Piemonte, Frataxin deficiency leads to reduced expression and impaired translocation of NF-E2-related factor (Nrf2) in cultured motor neurons, Int J Mol Sci 14 (2013) 7853–7865. 10.3390/IJMS14047853.

[53] R. Abeti, A. Baccaro, N. Esteras, P. Giunti, Novel Nrf2-inducer prevents mitochondrial defects and oxidative stress in Friedreich’s Ataxia models, Front Cell Neurosci 12 (2018) 188–198. 10.3389/fncel.2018.00188.

[54] D.R. Lynch, J. Farmer, L. Hauser, I.A. Blair, Q.Q. Wang, C. Mesaros, N. Snyder, S. Boesch, M. Chin, M.B. Delatycki, P. Giunti, A. Goldsberry, C. Hoyle, M.G. McBride, W. Nachbauer, M. O’Grady, S. Perlman, S.H. Subramony, G.R. Wilmot, T. Zesiewicz, C. Meyer, Safety, pharmacodynamics, and potential benefit of omaveloxolone in Friedreich ataxia, Ann Clin Transl Neurol 6 (2018) 15–26. 10.1002/ACN3.660.

[55] K. Itoh, N. Wakabayashi, Y. Katoh, T. Ishii, K. Igarashi, J.D. Engel, M. Yamamoto, Keap1 represses nuclear activation of antioxidant responsive elements by Nrf2 through binding to the amino-terminal Neh2 domain, Genes Dev 13 (1999) 76– 86. 10.1101/GAD.13.1.76.

[56] C.M. Clements, R.S. McNally, B.J. Conti, T.W. Mak, J.P.Y. Ting, DJ-1, a cancer- and Parkinson’s disease-associated protein, stabilizes the antioxidant transcriptional master regulator Nrf2, Proc Natl Acad Sci U S A 103 (2006) 15091– 15096. 10.1073/PNAS.0607260103.

[57] S.H. Choi, Y.W. Kim, S.G. Kim, AMPK-mediated GSK3beta inhibition by isoliquiritigenin contributes to protecting mitochondria against iron-catalyzed oxidative stress, Biochem Pharmacol 79 (2010) 1352–1362. 10.1016/J.BCP.2009.12.011.

[58] S.A. Hawley, M. Davison, A. Woods, S.P. Davies, R.K. Beri, D. Carling, D.G. Hardie, Characterization of the AMP-activated protein kinase kinase from rat liver and identification of threonine 172 as the major site at which it phosphorylates AMP-activated protein kinase, Journal of Biological Chemistry 271 (1996) 27879– 27887. 10.1074/JBC.271.44.27879.

[59] A. Woods, D. Vertommen, D. Neumann, R. Türk, J. Bayliss, U. Schlattner, T. Wallimannll, D. Carling, M.H. Rider, Identification of phosphorylation sites in AMP-activated protein kinase (AMPK) for upstream AMPK kinases and study of their roles by site-directed mutagenesis, J Biol Chem 278 (2003) 28434–28442. 10.1074/JBC.M303946200.

[60] D. Carling, The AMP-activated protein kinase cascade--a unifying system for energy control, Trends Biochem Sci 29 (2004) 18–24. 10.1016/J.TIBS.2003.11.005.

[61] M.C. Towler, D.G. Hardie, AMP-activated protein kinase in metabolic control and insulin signaling, Circ Res 100 (2007) 328–341. 10.1161/01.RES.0000256090.42690.05.

[62] M.S. Joo, W.D. Kim, K.Y. Lee, J.H. Kim, J.H. Koo, S.G. Kim, AMPK facilitates nuclear accumulation of Nrf2 by phosphorylating at Serine 550, Mol Cell Biol 36 (2016) 1931–1942. 10.1128/MCB.00118-16.

[63] V.B. Pillai, N.R. Sundaresan, G. Kim, M. Gupta, S.B. Rajamohan, J.B. Pillai, S. Samant, P. V. Ravindra, A. Isbatan, M.P. Gupta, Exogenous NAD Blocks Cardiac Hypertrophic Response via Activation of the SIRT3-LKB1-AMP-activated Kinase Pathway, Journal of Biological Chemistry 285 (2010) 3133–3144. 10.1074/JBC.M109.077271.

